# Improving confidence of differential transcription calls in enhancers

**DOI:** 10.1101/2025.09.12.675852

**Authors:** Hope A. Townsend, Jacob T. Stanley, Mary A. Allen, Robin D. Dowell

## Abstract

**Motivation:** Most disease-associated genetic variants reside within transcribed regulatory elements (tREs). Patterns of differential transcription at tREs can be leveraged to identify upstream regulators and link enhancers to their target genes. But the low transcription levels and high variability in tREs makes identifying high confidence differentially transcribed elements challenging.

**Results:** We present Mu Counts and TFEA-LE, two algorithms for robust identification of differentially transcribed tREs. The first step in accurate identification of differentially transcribed tREs is to obtain accurate RNA lengths and therefore counts over these regions. To this end we developed a method of accurate length inference (LIET-EMG) as wll as a rapid method for counting reads over tREs (Mu Counts). Armed with newly identified and quantified tREs, TFEA-LE then integrates motif information to simultaneously identify responsive tREs and their likely upstream regulators. We show improved precision and recall over general-purpose tools (e.g. DESeq2) in detecting p53-responsive tREs. We then clarify TF-specific responses within multi-TF perturbations in lung cells. Finally we show that the TFEA-LE approach improves TF activity inference, including in complex perturbations where many TFs respond. TFEALE is especially effective in technically challenging datasets, whether due to highly specific or broad responses, outliers, or high GC content. Ultimately, these 1 methods advance the systematic characterization of individual tREs, enabling their integration with regulators, target genes, and disease-associated variants for translational research.

**Availability and Implementation:** TFEA-LE: https://github.com/Dowell-Lab/TFEA/tree/Lead_edge. Nextflow pipeline to run Mu Counts: https://github.com/Dowell-Lab/Bidir_Counting_Analysis. LIET (including modifications for tREs): https://github.com/Dowell-Lab/LIET/tree/LIET_EMGtoo. Source code for this work: https://github.com/Dowell-Lab/Improving_tRE_Analysis_Paper

## 1 Introduction

Enhancers are genetic sequences that regulate genes—thereby allowing coordinated cellular responses and unique transcriptional states. Recent work on identifying enhancers has demonstrated that most enhancers produce lowly transcribed RNAs, hence they are more generally referred to as transcribed regulatory elements (tREs)[1]. These enhancer-associated RNAs have been found to be more reliable markers of local regulatory activity than epigenetic markers (e.g. H3K27ac ChIP-seq) [1–3]. While high-throughput measurements of nascent transcription, such as precision run-on sequencing (PRO-seq), accurately capture the transcription of enhancers, the tissue and perturbation specificity of enhancers leads to high biological variability between samples [2–4].

Measurements of enhancer transcription can vary significantly not only due to biological factors, but also technical artifacts such as depth, protocol, and analysis choices [1, 5]. This high variability complicates both the identification of tREs and detection of when they are changing [6]. Consequently, most efforts to date have focused on using tRE meta-profiles (rather than individual tREs) across conditions or samples to infer upstream regulators[7–9] or to link enhancers to their target genes[2, 10]. In these scenarios, general trends are detectable even if some changing enhancer RNAs are missed.

However, genome-wide association studies are revealing a growing need to confidently characterize individual tREs. The majority of disease-associated single-nucleotide polymorphisms (SNPs) fall within enhancers [11–13]. Given many SNPs can be inherited together (reside within linkage disequilibrium), functional data provides critical information for identifying specific causal variants[14, 15]. The relatively small size of tREs and their condition and tissue specific activities aid in fine-mapping of associated SNPs. This is particularly true when the tRE and its target gene both respond to a disease relevant perturbation[15]. These coordinated changes effectively pinpoint both the functional SNP and its relevant target, thereby implicating biological pathways, relevant cell types, genes, and/or upstream regulators. Consequently, annotating SNPs within enhancers holds tremendous potential to aid in the identification of potential drug-targets [11–13].

Yet identifying cell type or condition-specific tREs requires precise detection of changes in individual tREs. Although in-vitro methods like massively parallel reporter assays provide a method for high-throughput evaluation of tRE activity, they do not represent *in-vivo* conditions and can lead to inaccurate indication of SNP relevance and biological mechanisms [16]. Consequently, we sought to develop a pipeline for robustly identifying differentially transcribed tREs from high-throughput nascent run-on sequencing and similar enhancer-focused sequencing data (e.g. ATAC-seq). Our method builds on prior work on transcription factor enrichment analysis (TFEA)[7], leveraging both transcription changes (via ranking metrics) and motif information to support the identification of high confidence changes in tRE expression. With this approach, we simultaneously improve identification of responsive tREs as well as the upstream regulators that activate them. Additionally, we provide the first high-throughput length estimation of tRE RNAs, both enabling more accurate differential transcription measurement and facilitating future SNP integration. Overall, this work provides key steps toward incorporating individual tRE responses into downstream studies of biological mechanisms or SNPs for translational research.

## 2 Motivation

### Differentially transcribed tREs are poorly called by classic methods

Detecting differential transcription of individual tREs is becoming imperative for integrating noncoding SNPs and regulatory networks into drug target discovery. However, this advance has been largely stymied due to low confidence in identifying which tREs are significantly differentially-transcribed. Therefore, we first sought to evaluate how well current approaches detect differentially-transcribed tREs from nascent run-on RNA-sequencing data. However, to accomplish this goal, we need a reliable truth set—a set of tREs independently known to respond in a particular condition. Although no ground truth is known, we can take advantage of the fact that Nutlin-3a is a highly specific activator of transcription factor (TF) p53 and has been characterized by both nascent run-on RNA-sequencing and chromatin immunoprecipitation (ChIP) in multiple cell lines[17, 18]. For initial analysis, true p53-responsive tREs were defined by ChIP-seq peaks, while nonresponsive tREs required the lack of both a p53 motif and substantial ChIP-seq reads (details in Supplemental Methods). Using this truth set, we compared the three most commonly used differential expression tools (DESeq2, Limma, and EdgeR) under 13 parameter combinations (details in Supplemental Methods) to identify tREs responding to p53 activation[6, 19, 20].

Regardless of the cell type considered, recall of tREs with ChIP-seq peaks was poor (*<* 0.1) for all tools/parameter-combinations (Figure 1A and Supplemental Figure S1A). We wondered whether this performance was better than random guessing, so we defined an expected recall (based on chance alone, e.g. a “random set”) where tREs with increased transcription (positive log fold change) were randomly selected as responsive. As expected given the stringency of the negative truth set, very few, if any, non-responsive tREs (without motif or ChIP peak), were called significant except when using the random set (Figure 1A). Poor recall was consistent regardless of whether truth sets were defined by ChIP-seq peaks alone or both ChIP-seq peaks and motifs, and regardless of window size used for counting (Supplemental Figure S1A). As ChIP peaks do not inherently equate with transcriptionally responsive tREs and not all bona fide ChIP binding sites will have high quality motifs, we remained concerned that the poor performance simply reflected a reliance on ChIP for the truth set.

**Fig. 1.**
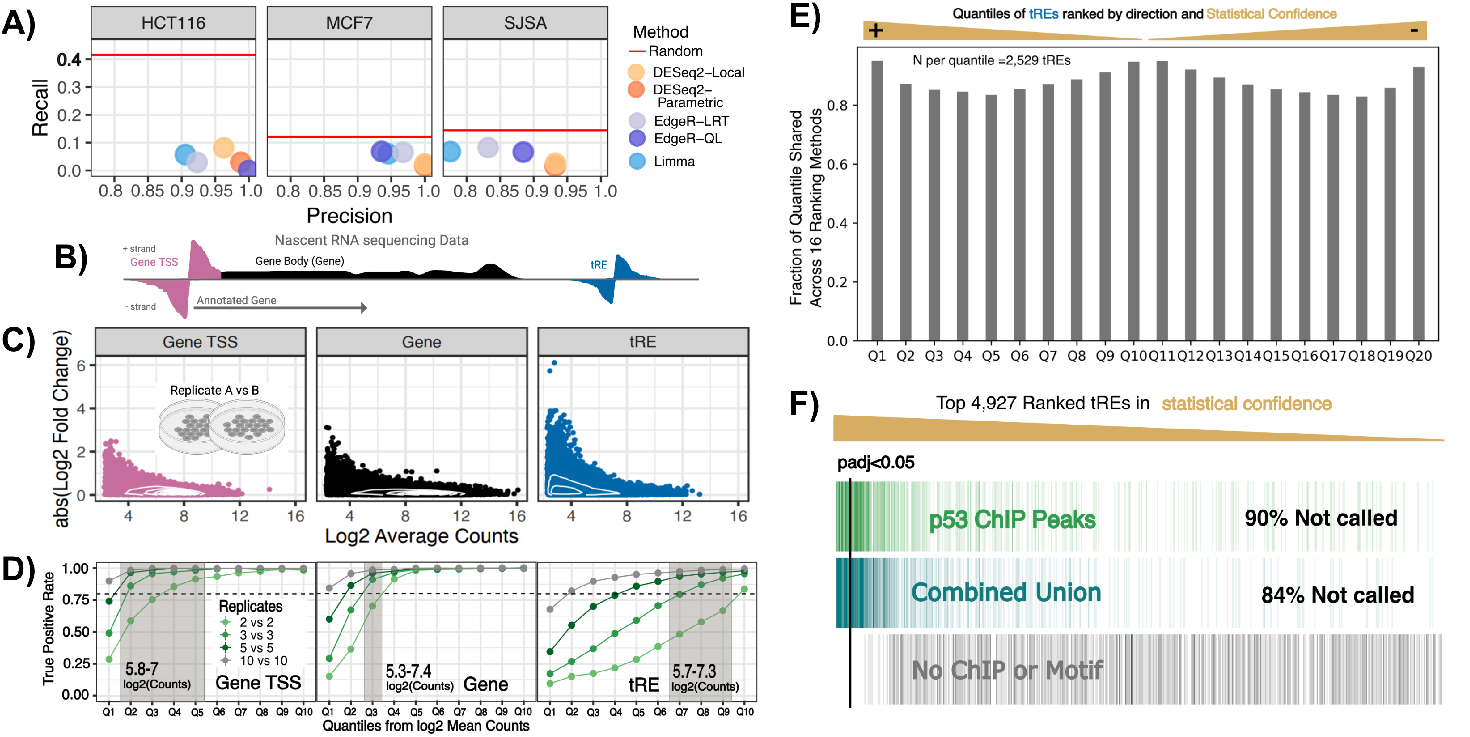
Despite classic calls of significantly changing tREs being poor, ranking based on statistical confidence is consistent. A. Recall and precision of p53 responsive tREs in multiple cell lines. Red line is recall from randomly assigning tREs with positive fold change as true. Truth based on p53 ChIP peaks and false positives lack motif or ChIP peak. Data from [18]. **B**. Cartoon representation of nascent run-on data at features including gene TSS bidirectionals (pink), gene bodies (black) and a tRE (blue) regions. **C**. Scatter and 2D density plots of read counts between biological replicates at various genome features. X-axis: log_2_ average counts, Y-axis: absolute value of log_2_ fold change. **D**. True positive rates (Y-axis) for genomic features across increasing replicate numbers (shades green) and quantiles (Q1-Q10) according to average counts (log_2_). Grey box represents regions with comparable average counts (≈5.5-7). **E**. The percentage of tREs within the same signed, ranked quantile according to all tested tool-parameter combinations except those using the Mean-based dispersion estimation from DESeq2 (see Supplemental Note). Ranking by adjusted p-value and direction of fold change. **F**. The top 6,479 tREs with positive log fold change after Nutlin-3a in MCF7 cells, ranked according to p-adjusted values. Green: overlaps p53 ChIP-seq peak, Blue: region called differentially transcribed when using all cell types as replicates (“Combined Union”), Grey: Lacking in p53 motif or ChIP peak. Black line: adj p-value cutoff of 0.05 for DESeq2-Local-LRT.

Activation with Nutlin-3a is a uniquely rich dataset, with multiple high quality replicates with high depth for each of three datasets (HCT116, MCF7, SJSA)[2, 18]. Therefore we next sought to evaluate a truth set based on only transcription. To this end, we considered the three cell types as replicates (six total replicates). Use of this combined dataset resulted in two truth sets, which include tREs called significant by any (“Combined Union”) or all (“Combined Intersection”) tools and parameter combinations (details in Supplemental Methods). When using these truth sets, we still see low recall, with a maximum of 60% of Combined Intersection and 30% of Combined Union called significant (Supplemental Figure S1A). Unrealistically large p-value cutoffs were needed to reach the recall enabled from using the random set, albeit at the cost of low precision (Supplemental Figures S1B). Indeed, poor capture of truth sets based on F1-scores are observed regardless of cell-type, tool-parameter combination, and truth set used (Supplemental Figure S2).

We predicted that the highly variable and low transcription of tREs[2] make them particularly challenging to call as differentially transcribed with high confidence. As a frame of reference, we compared the dispersion of tRE transcription to that of gene bodies and gene TSS associated bidirectionals. As expected, dispersion estimation (as measured by log 2 fold change) between replicates was highest for tREs when compared to gene bodies or gene TSS associated bidirectionals (Figure 1B). Notably, gene TSS associated bidirectionals – which are comparable in length to tREs – have lower dispersion of *log*_2_ fold changes than tREs between replicates, reflective of the fact that the majority of gene TSS associated bidirectionals contain higher counts than most tREs (Figure 1C). Thus, we next estimated the number of counts necessary to bring tREs to count levels comparable to gene TSS bidirectionals. To this end, we estimated that roughly a 2.5X increase (to about 100 million) in uniquely mapped and de-duplicated reads were minimally necessary (Supplemental Figure S3). An alternative to increased counts would be increased replication. To explore the impact of replication, we randomly sub-sampled reads from two high-depth samples. This simulation indicates that the vast majority of genes reach a true positive rate of 0.8 and false discovery rates close to 0.05 even with only two replicates of decent depth (40 million reads) (Figure 1D and Supplemental Figure S4B). While about 20% of genes and gene TSS associated bidirectionals are below the count threshold needed for this confidence, 70% of tREs are below it. Instead, around ten high quality replicates would be needed for tREs to reach similar confidence levels as what genes and gene TSS bidirectionals achieve with three replicates (Figure 1D and Supplemental Figure S4B). Consequently, based on our findings, extremely high depth and replicate numbers, not currently used across any known nascent run-on RNA sequencing experiments, are required to allow classic tools to reach statistical confidence in tREs comparable to genes [2].

Importantly, despite the poor recall and variable calls between the classic tools (Deseq2, EdgeR, and Limma) when using a single statistical cutoff, rankings of tREs based on direction and statistical confidence of change (p-value) are highly consistent (Figure 1E and Supplemental Figure S5). Indeed, the truth sets, such as p53 ChIP peaks and “Combined Union”, are enriched in tREs with the highest rankings (i.e. statistical confidence) (Figure 1F and Supplemental Figure S6). However, common significance cutoffs (e.g. alpha=0.05) of many tested parameter-tool combinations do not capture this enrichment well. This is especially true for the most popular tool according to citation numbers – DESeq2 – and across all tools in HCT116 (1F). One of the HCT116 samples has less overall transcription captured, and might therefore represent a more difficult dataset to analyze [2, 17]. Conversely, some parameter-tool combinations seemed to capture the enrichment of truth sets for MCF7 and SJSA samples well (Supplemental Figure S6). However, these approaches are the most permissive, and are therefore subject to extensive over-calling in datasets with more robust transcriptional responses[21, 22]. Additionally, much of the tREs supported to be changing from multiomics data have weak statistical confidence from transcriptional data alone. Therefore, we sought to develop a method to be robust to technical limitations by 1) leveraging the ability of classic statistical tools to effectively rank tREs on statistical confidence, and 2) integrating orthogonal biological data with this ranking to clarify true positives with low statistical confidence.

## 3 Algorithms and Pipeline

### A novel and reusable pipeline incorporates sequence-based support for calling tREs with poor statistical confidence

As high depth and replication are usually cost prohibitive, we sought a method for augmenting confidence by 1) maximizing transcriptional information gathered from nascent RNA-sequencing, and 2) integrating axillary biologically meaningful information.

An imperative first step in sequencing analyses is to identify the regions of interest, in our case, sites of unannotated bidirectional transcription (Figure 2A Step 1). In the original TFEA paper, tREs were identified using Tfit[23] per replicate and then combined across replicates into consensus regions for which differential transcription can be calculated. Because the position of RNA polymerase II initiation is critical to interpreting TF motif enrichment, *muMerge* was developed to maintain the best estimate for the midpoint of each replicate (which is assumed to be the position of RNA polymerase II initiation). To this end, *muMerge* assumes the width of the input region (e.g. called by Tfit) represents a confidence interval on the midpoint of the region, as this matches with Tfit’s output[23] (Figure 2A Step 2). This assumption results in a statistically principled method of obtaining high confidence estimates for the position of RNA polymerase II initiation across replicates – necessary for accurate TF activity inference[7]. However, as more replicates become available and confidence increases, *muMerge* has the unintended consequence of outputting region widths that no longer reflect the transcribed expanse of the replicate data[2]. Because TFEA counts over these *muMerge* output regions, we realized the consensus regions were under-estimating the transcript levels measured within the replicates. Consequently, for counting purposes, a fixed window around the center point was previously used[2] (Figure 2A Step 3). This approach, however, does not take noise from overlapping transcripts into account, which is commonplace in nascent run-on sequencing data.

**Fig. 2.**
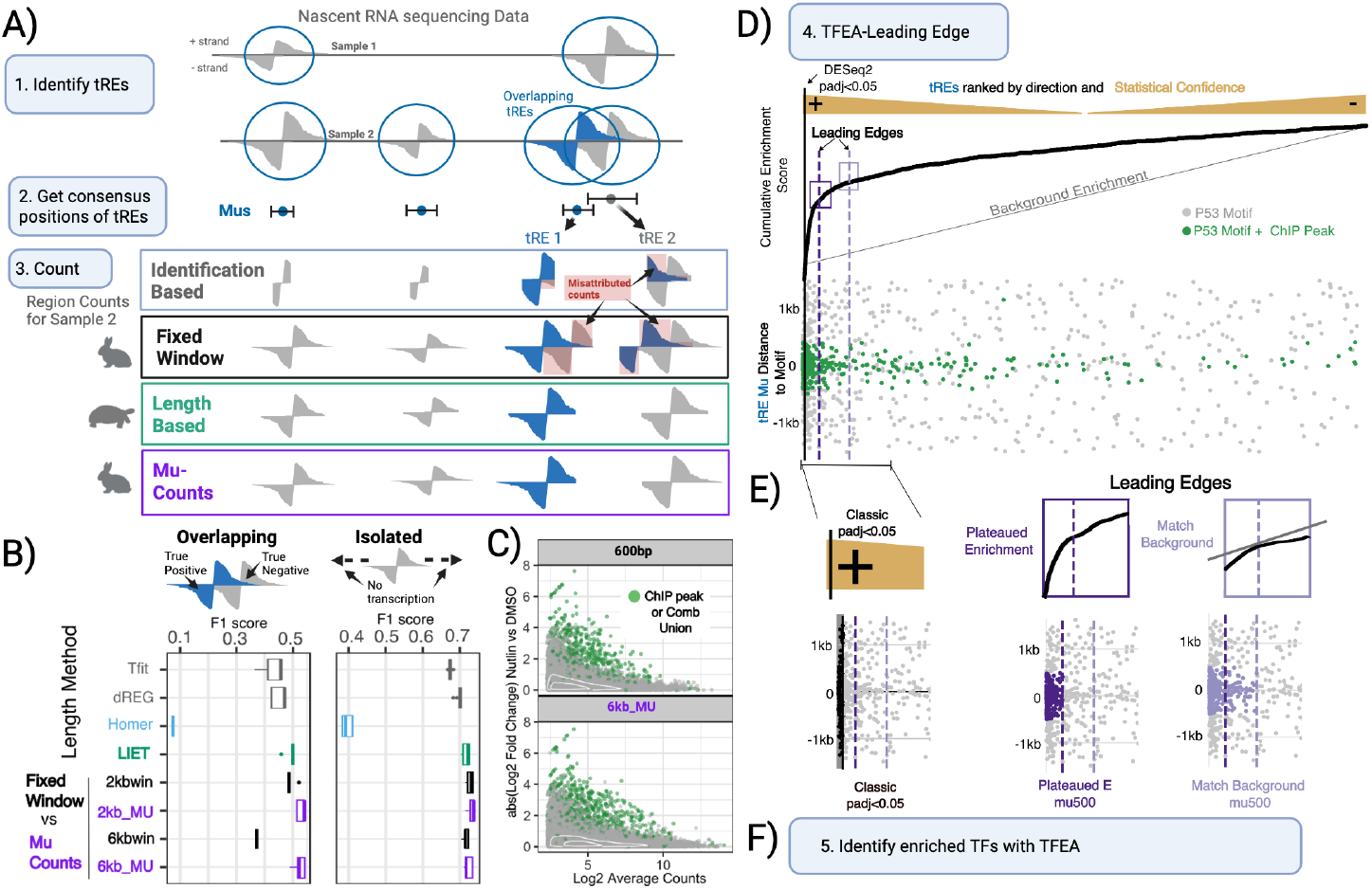
An improved workflow for identifying differentially transcribed tREs from nascent run-on sequencing data. A. First three steps involve 1) identifying tREs, 2) identifying consensus regions from multiple replicates, for which we show muMerge[7], and 3) counting reads over consensus regions. Shown are four methods of counting: Identification based (using Tfit or dREG), Fixed window, Length based (in this case, LIET), and Mu Counts. Misattributed counts (red highlight) indicates counts that would be falsely considered for one tRE despite belonging to another. Fixed Window and Mu Counts are both fast (rabbit) while LIET is slower (tortoise). **B**. F1 scores (harmonic mean of recall and precision) for p53-responsive tREs when using lengths according to various methods: tRE identification tools (Tfit and dREG, grey), HOMER (blue), LIET (green), fixed windows (black), and Mu Counts (purple). Isolated tREs have no detectable transcription (tRE or gene) within 8kb. **C**. Scatter plots of *log*_2_ fold change between Nutlin-3a and DMSO cells of tREs when using counts based on 600bp fixed window (top) or 6kb Mu Counts (bottom). Truth sets according to ChIP-peaks or “Combined Union” are in green. **D**. Once regions are counted and ranked (top) by direction and significance, the leading edge is calculated on the enrichment curve (middle). The enrichment score reflects co-occurrence with motif instances (bottom, dots), with each motif instance weighted by the distance of the motif to tRE center (*µ*, labeled 0). tREs with p53 ChIP support are colored green. **E**. Two methods of leading edge detection are compared to the classical statistical cutoff from DESeq2 (left). Plateaued Enrichment considers when the cumulative enrichment score growth stalls significantly (dark purple). Match Background identifies when the slope of the enrichment curve matches the expected background (light purple). Final leading-edge calls can consider tREs with motifs within a certain distance of *µ* and within the leading edge (e.g. mu500 = 500bp, full=All). **F**. Finally, TFEA enrichment scores are used, with optional GC-bias correction (TFEA) or leading edge adjustments, to identify responding TFs.

To address this counting problem, we revisited the numerous tRE identification algorithms and ask how accurately the region calls reflect the transcript length. Because tREs are primarily unannotated, they must first be identified by one of the numerous tRE identification algorithms[24]. Prior work benchmarked the ability of tools to identify active tREs, finding Tfit was the most accurate at capturing the midpoint of the bidirectional region (i.e. the position of RNA polymerase initiation)[25]. However, the benchmark did not ask whether the lengths of tREs accurately reflected their transcribed length. Therefore, we compared the output length from a subset of these algorithms (Tfit, dREG[26], Homer[27]) to lengths determined by previously published long-read sequencing in the same cell type [28–30]. Importantly, while Tfit and dREG were built to specifically identify bidirectional regions, Homer was built to capture transcript regions on each strand, independently. Additionally, we include a modified version of the recently published LIET approach[31] that focuses explicitly on capturing transcription lengths of bidirectional regions (details in Supplemental Methods). Unlike Homer, LIET can incorporate the consensus midpoint positions from *muMerge* for final coordinates, outputing both the tRE midpoints and endpoints. When applying these approaches to transcriptionally isolated tREs, we found that identification methods (e.g. Tfit and dREG) tended to largely underestimate length estimates while Homer and LIET lengths were closer to those suggested by long-read sequencing (S7A). Most tREs, however, are found within genes (e.g. intronic) or overlap other tREs [2] (2A). Therefore, we also simulated high noise (random reads distributed) to represent overlapping transcription. While Homer falsely captured noise as extensions of transcripts, LIET-based estimates were unchanged (S7B). Instead, the LIET model’s background parameter effectively and accurately captured the introduced noise and gave predicted lengths closest to the long-read data (Supplemental Figure S7B+C). Despite the accuracy of the LIET approach, the underlying algorithm is inherently computationally expensive and can quickly become obtuse when considering thousands of tREs (20,000 tREs in one sample took *>* 24 hours) (Supplemental Figure S7D).

Given that our overall goal is to maximize the signal of tREs while avoiding noise from overlapping transcription, we wondered if a faster method could be developed to obtain optimal counts per tRE, without requiring apriori knowledge of accurate RNA lengths. This approach would ideally incorporate *muMerge* consensus midpoints. To this end, we developed a pipeline (Mu Counts) that seeks to rapidly identify reads in a fixed window (around consensus midpoints as from *muMerge*) that can be definitively assigned to a specific tRE, even when another tRE is nearby (Figure 2A Step 3 “Mu-Counts”). A figure describing the full pipeline can be found in Supplemental Figure S8. This approach is fast, able to consider all relevant samples in a 15 minute time frame (Supplemental Figure S7D). We next asked to what extent the improved counts would, by themselves, improve our ability to call differentially transcribed tREs. Both LIET and Mu Counts not only improved differential calls of transcriptionally isolated tREs, but successfully distinguished between overlapping tREs with and without multi-omic evidence of change (“Overlapping”) (Figure 2B). Mu Counts and LIET’s having high and comparable rankings were consistent across all cell types (Supplemental Figure S9). Importantly, Homer’s low F1 scores arise in large part because it fails to call many regions (Supplemental Figure S10). Instead, LIET and Mu Counts can focus on tRE characterization based on accurate consensus midpoints of tREs (e.g. from *muMerge*) [7].

Despite effectively maximizing transcriptional data retrieval, we still observe low statistical confidence for most tREs. While Mu Counts indeed shifts the density of mean counts, dispersion trends don’t improve; instead many expected true positives still show transcriptional responses difficult to distinguish from noise(Supplemental Figure S11 and Figure 2C). Therefore, we still sought to improve identification of true positives that have limited data from transcription alone.

One way to improve our ability to determine which tREs are differently expressed would be to incorporate additional information. Perturbations that affect tREs transcription do so through the alteration of transcription factor function. Moreover, transcription factors most often work through DNA motifs. Therefore, we wondered if information surrounding the motifs within the tREs could be used to further improve our ability to detect differently transcribed tREs. Given the clear correspondence between the tREs with multi-omics support and tREs with the highest ranking for statistical confidence (Figure 1E), we sought to develop a rank-based method of identifying differentially transcribed tREs that could integrate such additional data. For this, we considered how we have already shown the power of ranking and motifs in TFEA, which uses motif co-occurrence with sites of bidirectional transcription (at tREs or gene TSS) to robustly quantify changes in TF activity between conditions[7, 8, 32]. In TFEA, tREs are first separated by direction of fold change and then ranked by the differential p-value as defined by tools like DESeq2. Iterating through the ranked tREs, TFEA then calculates a cumulative enrichment score, with the score increasing according to the proximity of a specific TF motif to the tRE’s midpoint [7]. When TF motif instances co-occur with the tRE midpoint at the extremes of the ranking, the TF is inferred as actively contributing to the differences between the conditions (Figure 2D). Therefore, we hypothesized that identifying the tREs most contributing to a given TF’s significant call in a perturbation would correspond well with tREs responsive to a perturbation. This concept is similar to gene set enrichment analysis identifying a “leading-edge” to pinpoint which genes are responsible for a pathway’s call, and therefore relevant to the perturbation[33]. Consequently, we developed a similar approach, instead identifying the “leading-edge” of the enrichment curve within TFEA. Recall that the cumulative enrichment curve is calculated as a weighted distance of a TF motif instance from the tRE center. Consequently, the inflection point of this curve represents a statistically principled shift in the enrichment of TF-motif instances and changes in transcription (Figure 2D + E) and hence would be an alternative metric for defining regions of differential transcription that utilizes both transcription and sequence signals.

To identify the inflection point (aka leading-edge) we developed two approaches, one defines a conservative inflection point for identifying changing tREs, and the other captures large scale trends of enrichment (Figure 2E). Specifically, the first looks for the point where the rate of change in the enrichment curve plateaus, hereafter called “Plateaued E” where E stands for Enrichment. The second identifies the point at which the slope of the cumulative enrichment curve is comparable to the TF motif presence in unchanged (background) tREs, hereafter called “Match Background”. Details regarding these approaches can be found in Supplemental Methods. Both algorithms for obtaining the leading-edge (Plateaued E and Match Background) are largely consistent regardless of the classic statistical tool-parameter combination used to rank tREs (Supplemental Figure S12). This approach extends beyond relying on a single p-value cutoff with classic statistical tools. Instead, a user can consider tREs based on multiple sources of confidence–both sequence support and statistical analyses for transcriptional change–to assess downstream biological questions.

Finally, analyses usually end with identifying the TFs that are active within a perturbation or condition, as with TFEA (2F Step 5). Sequence-based predictions, however, are hindered by the fact that transcription initiation sites (including tRE midpoints) tend to be GC-enriched[8, 32]. TFEA attempts to mitigate this bias by using a heuristic correction term to enrichment scores before deciding significance based on a simple regression model[7]. This was admitted to be a rudimentary solution in the original paper, and so we end this work with evaluating how to improve upon it using what is learned from the leading-edge [7].

Overall, the resulting pipeline (Figure 2) still takes advantage of consensus positions from *muMerge* before combining improved feature counting (Mu Counts) with the leading-edge approach of TFEA, the latter of which is hereafter referred to as TFEA-LE.

## Implementation and Results

### TFEA-LE improves understanding of single-TF and multi-TF transcriptional responses

We sought to test TFEA-LE, first by comparing it’s calls to the regions identified by the padj *<* 0.05 cutoffs of classic tools (both using Mu Counts). We find that considering tREs within either leading-edge algorithm significantly improves recall of p53 ChIP-peaks, while maintaining high precision 3A. These improvements, as also measured by F1-scores, are found in all cell types, regardless of how we define truth (ChIP-peaks, Combined Intersection, Combined Union), parameter-tool combination, or length of windows used for Mu Counts (Supplemental Figure S13). Figure 3B shows an example of a tRE that, despite having p53 and Nutlin-specific H3K27ac peaks across all cell types, would only be considered significantly changing in response to Nutlin-3a in HCT116 cells from classic statistical approaches. Using Mu Counts allowed MCF7 to indicate a significant call as well, compared to using fixed window counts. Only the leading-edge (TFEA-LE) allowed this tRE to be called significant across all three cell types, hence matching the multiomics support. tREs only called by TFEA-LE also had increased read counts in Nutlin-3a (p53 activating condition) compared to control, more so with Plateaued E (Supplemental Figure S14). Likewise, the leadingedge calls are highly enriched in previously published Nutlin-induced H3K27ac peaks in MCF7 and HCT116, more so than tREs called from classic statistical approaches (SJSA H3K27ac data are not available) (Figure 3C and Supplemental Figure S15). The leading-edge identifies a much larger number of p53 responding tREs shared across all cell types compared to the original publication using classical statistical analysis methods[18] (Supplemental Figure S16A). Despite this, the percentage of tREs called shared across all cell types that are also supported by p53 ChIP peaks [shared across cell types] is consistently higher for LE-based calls than those from only classic tools (Figure 3D, Supplemental Figure S16B).

**Fig. 3.**
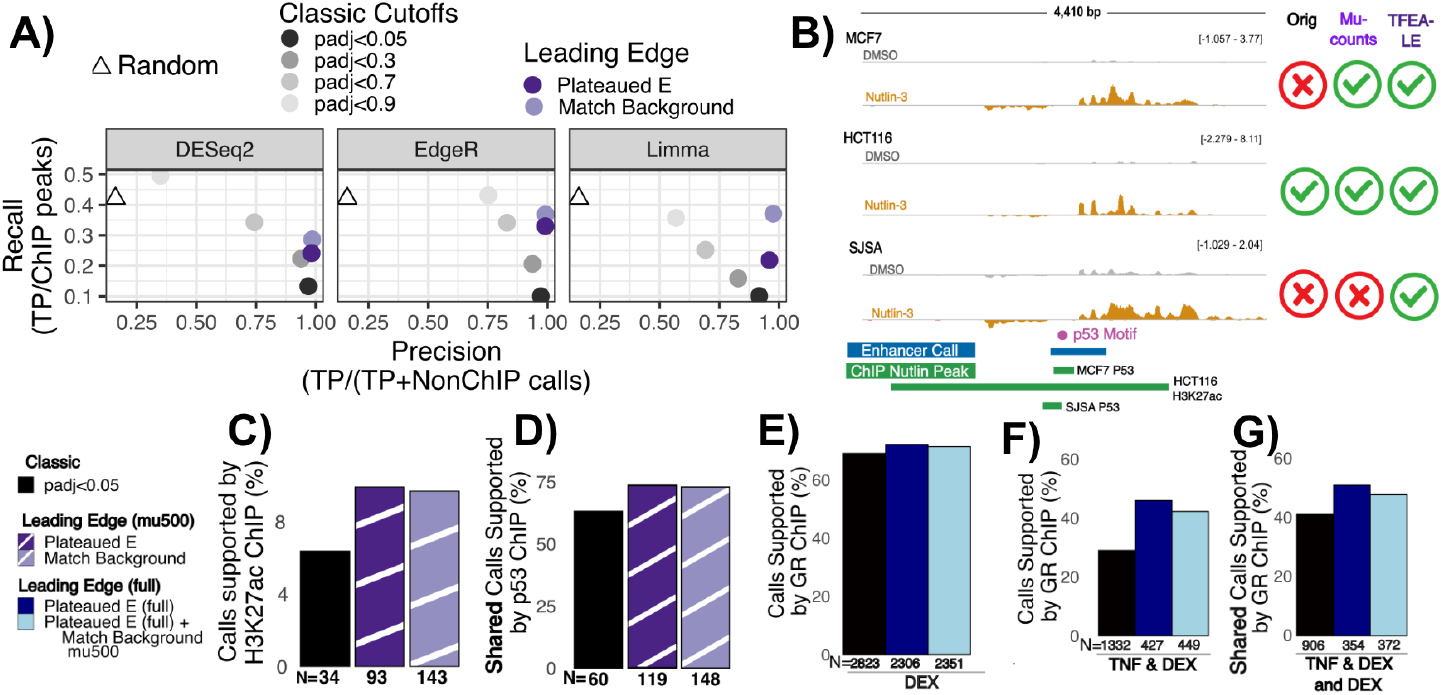
The leading-edge approach enables improved recall of Nutlin-responsive and DEX/TNF-responsive tREs across cell-type and perturbation combinations. A. Precision and Recall of HCT116 tREs with p53 ChIP peaks. Triangle: randomly selected tREs with positive fold change; Grey dots: adj. p-value cutoffs; Purple dots: Leading edge methods (classic padj *<* 0.05 + mu500). Methods correspond to DESeq2-Local-LRT (left), EdgeR-Locfit.mixed-QL (middle), and Limma-Trend-eBayes (right). **B**. An example region (chr2:113612107-113616515) for three cell types before (grey) and after Nultin-3a (orange). Data from [18]. Check-marks (far left) indicate whether the tRE was called significant in that cell type with different approaches. Org: 2kb fixed window; Mu Counts: 2kb; TFEA-LE ranked based on 2kb Mu Counts. **C**. Percentage of HCT116 Nutlin-responsive tRE calls from the different approaches that also have a H3K27ac peak only after Nutlin was added to the media. Black: padj*<* 0.05 by TMM-eBayes-Trend; Dark Purple: Plateaued E with padj*<*0.05 + mu500; Light Purple: Match Background with padj*<*0.05 + mu500. **D**. Percentage of calls shared across all cell types, with p53 ChIP in Nutlin-3a also shared across cell types, according to the different approaches. **E**. Percentage of tRE calls in DEX-treated cells that also have a GR ChIP peak. Black: DESeq2-Ratio-LRT-Local padj *<* 0.05; Dark Blue: Plateaued E (full); Light Blue: Plateaued E (full) + Match Background mu500. **F**. Percentage of tRE calls in DEX and TNF treated cells that also have a GR ChIP peak. Leading-edge results are based on the GR TF motif. **G**. Percentage of tRE calls shared between cells exposed to both TNF and DEX and just DEX that also have GR ChIP peaks shared between the two conditions. In all cases, Ns listed are the total number of tREs called significant by each approach. Method selected for black (C-G) was selected for highest number of shared tREs across all samples.

Arguably, Nutlin-3a is an exquisitely specific activator of a single transcription factor – p53. Thus, we next wondered whether the leading-edge approach works well when multiple transcription factors are responding to a perturbation. To assess this, we applied TFEA-LE to our previously published double drug study[34]. In that work, we exposed Beas-2B cells to either dexamethasome (DEX), TNF-*α* (TNF), or both drugs. DEX activates glucocorticoid-based receptors (GR and MCR) while TNF activates the NF*κ*B complex, including multiple transcription factors (REL, RELB, TF65, NF*κ*B1, NF*κ*B2) [34]. All calculated leading-edges for these TFs showed similar consistency across ranking methods, as previously found with p53. Indeed, the leading-edges also had greater consistency in the differential tREs called than classic statistical methods (Supplemental Figure S17). Like observed with Nutlin, the rankings of tREs with greatest statistical confidence across tools were consistent, further supporting the applicability of a rank-based approach (Supplemental Figure S18). Unlike with p53, however, classic statistical tools called more tREs significant than the Plateaued E leading-edge values indicated (Supplemental Figure S19). We predicted that the greater number of tREs called by (e.g. DESeq2 and EdgeR) could be due to these tools not being specific towards upstream regulators, while the leading-edge is TF-specific. These drugs also produce highly robust transcription compared to many other nascent sequencing experiments, suggesting that classic statistical tools are less disadvantaged in predicting tREs that change or might be prone to overcalling [2].

Therefore, in this case, we hypothesized that the leading-edge would allow 1) secondary support for which adjusted p-values should be used for classic tools in cases with robust transcription and 2) more specific identification of tREs being regulated by a single TF of interest. To test these hypotheses, we focused on considering all tREs within the leading-edge, instead of just those with strong motifs. This approach would still consider motif enrichment trends, which could be TF specific, but allows for cases where TF motifs aren’t as precise, considers secondary responses, and takes advantage of more robust transcriptional data. When applying this approach to the DEX condition (GR focused activation), we indeed observed either a slightly increased or comparable enrichment of calls supported by GR ChIP-seq with the leading-edge compared to classic statistical tools (Figure 3E and Supplemental Figure S20A).

To test our second hypothesis, that the leading-edge could identify tREs most likely regulated by a specific TF, we also assessed if we could clarify GR-specific tREs amidst conditions where multiple TFs were perturbed (DEX+TNF). When considering cells perturbed with both DEX+TNF, the GR leading-edge tREs were further enriched in GR ChIP peaks than those called by classic statistical approaches; this trend is observed even after removing tREs called significant in cells perturbed with TNF alone (Figure 3F and Supplemental Figure S20B). Next, we compared tREs responding with just DEX, and in cells perturbed with both TNF and DEX. Again, the leading-edge calls shared between these perturbations had a higher enrichment of ChIP calls also shared between the two perturbations, regardless of classic statistical approach used; leading edge calls for perturbation-unique tREs were also more enriched in ChIP-seq peaks unique to a perturbation 4G and Supplemental Figure S20B)).

### The leading-edge improves TFEA-based analyses of technically challenging multiomics data

In the case where a single TF is primarily responding (e.g. Nutlin-3a or DEX), a single motif is found in most differential tREs. However, in perturbations with multiple TFs responding, the motifs for a single TF are further apart in the rankings in the ranked tREs. Indeed, the more TFs that are responding, the more likely motifs of significant TFs will be found across a larger spread of ranked tREs rather than just those at the extreme poles. A larger fraction of tREs responding also introduces a larger number of TF motifs that can arbitrarily be enriched in tREs changing. This trend is particularly common for motifs with high GC content (due to the GC-enrichment of transcription-initiation sites), further exacerbating GC-bias. To mitigate this bias in TFEA, corrected Enrichment-scores are calculated as a y-offset of the observed Enrichment-scores from a linear regression fit (extreme case shown in Figure 4A). However, over time, we have found that this approach fails with strong GC bias, introducing as many false-positive TFs as it mediates against [7]. Therefore, we next sought to evaluate whether leading-edge metrics can also be used to reduce false-positive TF enrichment calls.

**Fig. 4.**
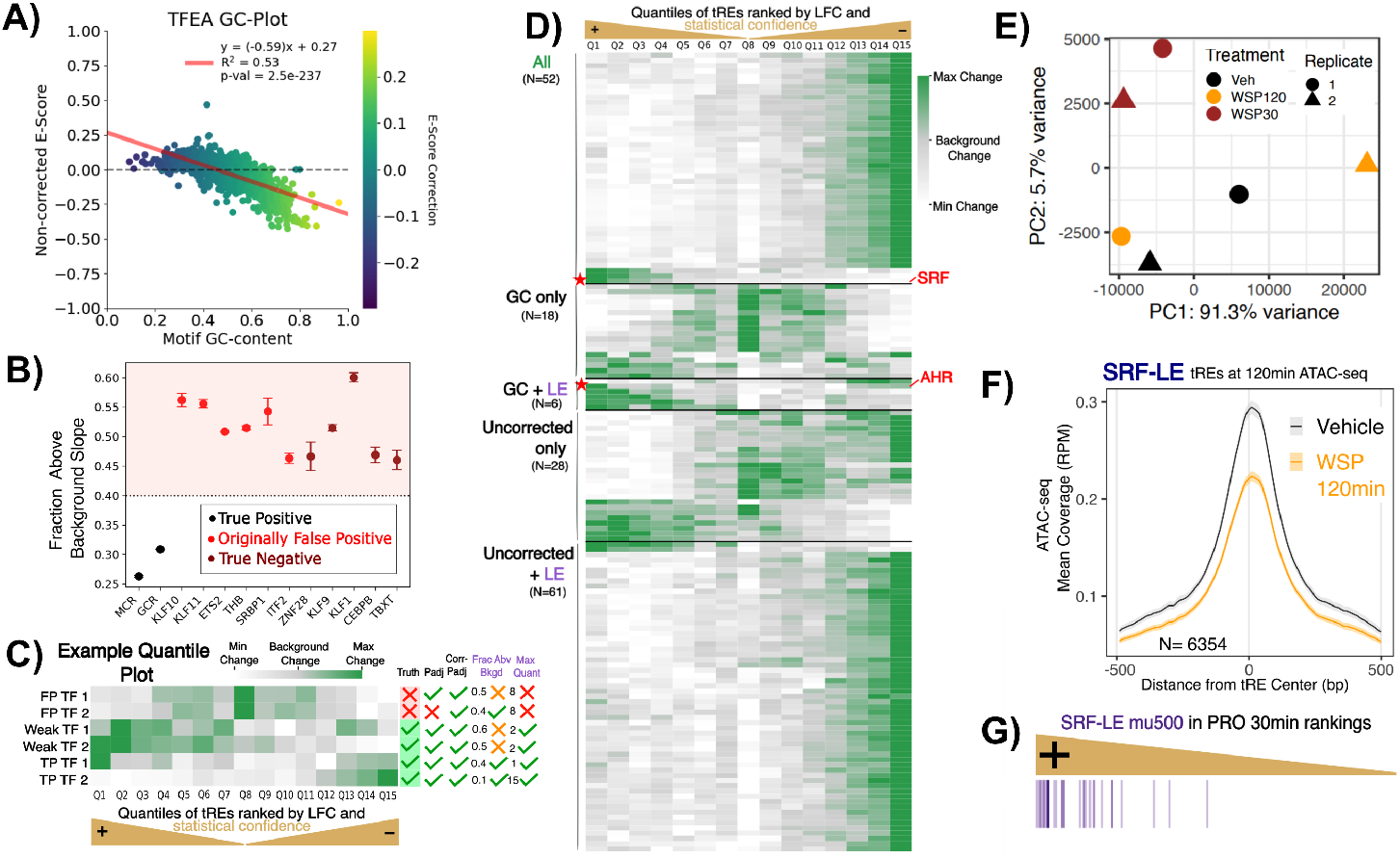
The Leading Edge improves identification of true TF responses and responding tREs in technically challenging multi-omics data. A. TFEA enrichment score plot for ATAC-seq at 120 min. Dots colored by GC-content, red line: regression fit. **B**. Fraction of tREs with cumulative enrichment scores above expectation from background (y-axis) is plotted for numerous TFs (x-axis). Dots are colored by correctness status based on TFEA enrichment score alone. The area about 0.4 is shaded in pink as a consistent cutoff between true positives and negatives. **C**. Cartoon example of quantiles within the ranked list, colored by their median change in enrichment. Coloring sets grey as expected background level (“background change”). Patterns represent two false positives (FP), two weak TF signals, and two strong TF signals (TP). Check marks on right indicate whether the TF is considered significant by different methods. TFEA’s Padj and GC-corrected (Corr-Padj), Frac. Abv. Bkgd (Fraction tREs with cumulative enrichment increase above background expectation), and Max Quant (requires maximum value at edges of ranked list). **D**. Quantile heatmap (similar to C) based on PRO-seq from BEAS-2B cells 30 minutes after their perturbation with wood smoke[35]. TFs (rows) are sorted by whether they are called by TFEA uncorrected, TFEA GC-corrected, or TFEA-LE methods. TFs with extensive wet-lab validation are marked with red stars (SRF and AHR). Change is calculated via a binning of ranked tREs (each bin N=2994). **E**. PCA plot of ATAC-seq BEAS-2B samples without WSP (Veh) or with WSP perturbation for 30 or 120 minutes (WSP30 and WSP120) and replicate 1 or 2. **F**. Metagene showing normalized ATAC-seq reads after 120 minutes for cells perturbed with vehicle (grey) or wood smoke particles (orange) of regions within the Plateaued E leading edge for Serum Response Factor (SRF). **G**. tREs within the Plateaued E leading edge for SRF at ATAC-seq 120 minutes (decreasing accessibility, with SRF motifs within 500bp) are colored as they are ranked by differential transcription confidence according to PRO-seq at the 30min mark (focusing only on tREs with positive log fold change).

In the Nutlin-3a and TNF/DEX data sets, we observed that false-positive TF calls and true negative TF calls by TFEA consistently had Match Background leading-edges that suggested either half or zero tREs were contributing to the TF’s enrichment score. Remarkably, these cases suggested that some significant enrichment scores were being called based on tREs with no detectable transcriptional change between conditions (Supplemental Figure S21 top). To robustly measure this trend, we calculated the fraction of tREs contributing to cumulative TF enrichment scores above what is expected from background noise (hereby called “Fraction Above Background”). Indeed, we saw that while DEX-induced transcription factors GR and MCR have lower fractions, false-positive and true negative TF calls have fractions above .45 (4B). This finding is reproducible across all Nutlin-3 and DEX/TNF perturbed cells, with the fractions being lowest for highly specific TF responses (e.g. P53) (Supplemental Figure S21).

Furthermore, we considered that TFEA uses a fixed motif scanning cutoff for all motifs, which results in some motifs having far more called instances due to their sequence being more likely by random chance alone. To address this bias, we enabled different p-value cutoffs for motif scanning to be used per transcription factor motif. Default cutoffs for each motif were optimized as a function of the number of motifs found across all tREs of the genome, resulting in different adjusted-pvalue cutoffs per motif (details in Supplemental Methods). This eliminated the gross over-calling inherent with some motifs.

As seen with the TNF/DEX data, as more TFs respond, the response to any single TF may not be perfectly clustered at the extremes of changing transcription. Instead, signals from multiple TFs may intermingle, effectively broadening the number of altered tREs and consuming more of the ranked list with responsive tREs. However, TFEA – by using the middle quartiles as background – assumes that the enrichment score would be contained exclusively within the first or fourth quartile. To ask whether this assumption holds for complex perturbations, we divide the data into fifteen equally sized bins (15-quantiles: Q1-Q15) and plot the maximum enrichment score per quantile (Figure 4C). We observe three general patterns for motifs: 1) strong motif enrichment at the extremes, as expected by TFEA; 2) weaker enrichment at the extremes, as is typical for weaker responders when multiple TFs respond; and 3) instances where the strongest motif enrichment is in the middle of the ranking, where tREs have little to no detectable change in transcription. This last group is likely false-positives. We predicted that using this positional enrichment assessment, we could both separate out weak and stronger responders, as well as remove false-positive TF calls.

Using these refined correction metrics, we turned our attention to complex perturbations where multiple transcription factors are responding. To this end, we re-examined paired PRO-seq and ATAC-seq data generated from Beas-2B cells perturbed with wood smoke particles (with two time points: 30 and 120 minutes) ([35]). This perturbation produces a very robust transcriptional response, with many TFs responding. These data also have a strong GC bias in their active tREs, leading to a large linear regression coefficient in traditional TFEA (Figure 4A and Supplemental Figure S22). The default GC-bias correction paradoxically results in artifically shifting tens of TFs into significance, most of which have no support in the literature for being involved in wood smoke or respiratory response. However, the GC-bias correction also enabled the calling of the TF Aryl Hydrocarbon Receptor (AhR), one of the best-known respiratory responders with confirmed activation in these same cells according to ChIP-qPCR, siRNA knockdown, and western-blots [35]. We wondered whether the improved TFEA-LE approach would both identify AhR and remove the large number of apparent false-positives.

To assess the value of this approach for the wood smoke particle response, we split TFs according to whether they were being called significant by TFEA’s GC-corrected scores, TFEA’s uncorrected scores, and/or TFEA-LE based metrics. For initial evaluation, we used strict TFEA-LE based metrics, requiring *<*0.45 “Fraction Above Background” and the quantile with the maximum enrichment score (“Maximum Quant”) outside the middle (max q *<* 5 or *>* 10). When examining PRO-seq 30 minutes, TFs supported by all methods showed very clear maximum changes in cumulative enrichment in tREs with the greatest confidence of statistical change (Figure 4D). This included the transcription factors first noted in the original publication for response: Serum Response Factor (SRF) [35]. The leading-edge approach also uniquely confirmed the significance of AhR (Figure 4D star). Conversely, most TFs called by GC-bias correction alone, and many called with uncorrected scores alone, had maximal enrichment scores in middle quantiles, consistent with false-positives (Figure 4D). These trends are generally visible at all time points and in both PRO-seq and ATAC-seq (Supplemental Figure S23). As a final support for calls, we considered which TFs were supported as relevant by both ATAC-seq and PRO-seq, in the same direction of enrichment. The percentages of TFs called with the same enrichment direction in both ATAC/PRO were highest in TFs shared across all calling approaches, and next LE-based calls at all time points (Supplemental Figure S24). A comparable but less consistent trend was observed for agreement in direction when using gene TSS bidirectionals compared to tREs (Supplemental Figure S24).

We next examined whether TFEA-LE would work equally well on the ATAC-seq data as on nascent run-on sequencing data since TFEA itself has been applied successfully to ATAC-seq [7]. In the wood smoke data, both DESeq2 and EdgeR called a maximum of one region significantly changing chromatin accessibility at the 120 minute mark, likely due to one of the two 120 minute samples showing outlier accessibility patterns, even after normalization (Figure 4E). Despite classic approaches indicating no tREs were changing, TFEA called several TF motifs as significantly enriched in the regions ranked towards decreasing accessibility at 120 minutes, including SRF. Therefore, we assessed if the leading-edge tREs of SRF would represent tREs changing in accessibility, despite generally weak agreement between samples. Indeed, normalized counts of the Plateaued E leading-edge regions for SRF show a clear depletion in accessibility at 120 minutes with wood smoke particles (Figure 4F). Additionally, while SRF-based leading-edge tREs showed decreased accessibility in both 120 minute replicates, random tREs of the same number that had fold changes below 0.9 only showed decreased accessibility in one replicate (Supplemental Figure S25). Finally, SRF leading-edge tREs with SRF motifs have increased transcription at the 30 minute mark (which follows SRF activity at that time point) before having transcription levels again comparable to Vehicle by 120 minutes (Figure 4G). Similar plots of 4F and G for all the time points and leading-edge approaches show similar trends and can be found at (https://github.com/Dowell-Lab/Improving_tRE_Analysis_Paper/WSP). Therefore, we successfully identified tREs with changing accessibility at 120 minutes as supported by other data, even when classic statistical cutoffs of original tools could not.

## Discussion

Overall, in this work we provide a new collective suite of tools to improve the characterization of transcriptional regulatory networks from multiomics-data. Although especially powerful when integrated together, all tools are available as independent software packages to best fit a user’s needs. The high dispersion and low transcription of tREs present a particularly difficult challenge for traditional statistical tools such as DESeq2. To identify high-confidence differentially tREs, we developed two new tools to augment the TFEA algorithm: Mu Counts and TFEA-LE. Mu Counts addresses original limitations of *muMerge* by maximizing the data we can consider from tREs through improved counting, while still taking advantage of consensus positions of *muMerge*. Additionally, TFEA-LE leverages motif co-localization signals to assist in the identification of statistically significant changes in transcription signal at lowly transcribed regulatory elements. For this purpose, TFEA-LE improves on the original TFEA algorithm in multiple ways. First, TFEA-LE uses a leading-edge approach to identifying the inflection point of co-enrichment of motif instances with the extremes of transcription changes. Second, cutoffs for motif scanning now reflect frequency of hits for that motif. Finally, we leverage the TFEA-LE metrics to improve discrimination between true positive enrichment and false-positive TF calls. Collectively, these improvements enable TFEA-LE to both identify differentially transcribed tREs that are supported by multiple lines of biological information and to address complex perturbations that can have a either a highly specific or very broad impact on cells.

While prior work has benchmarked the identification of tREs[1], we extend this work to the question of whether these methods accurately return the lengths of the transcribed region. In this extension, we also introduce a tRE-focused adaptation of the LIET model, finding that the LIET model’s explicit background parameterization provides a uniquely powerful approach for distinguishing noise from signal, resulting in the best length estimates. However, LIET is computationally expensive, and tools such as Homer present an effective alternative for tREs with confirmed isolation from neighboring and overlapping transcripts. Accurate lengths of enhancer-associated transcripts are necessary for characterizing the RNA itself as well as using them in fine-mapping of SNPs. Indeed, we and others have already shown that using tREs can be essential towards filtering and annotating SNPs[14, 15], but were confined to about 1% of SNPs as we focused on only the regulatory region driving the tRE’s transcription (i.e the initiation/loading zone[36]) rather than the RNA produced. Indeed, we now know that a subset of these transcripts can be functional in ways that tRE RNA SNPs can lead to disease-related phenotypes [37–41].

The introduction of Mu Counts was necessary to bolster counts associated with individual transcripts. The method is a fast heuristic that accounts for overlapping transcription of both genes and tREs, counting only regions where reads can be conclusively assigned. The accuracy of Mu Counts, however, is dependent on the accurate identification of tREs and their midpoint, i.e. the position of RNA polymerase II initiation to correctly disentangle overlapping tREs. Hence, the combination of *muMerge*[7] which seeks to preserve accuracy on the inferred midpoint of tREs across replicates, with Mu Counts results in a measurable improvement in differential-transcription analysis of tREs through increased recall. Beyond differential transcription analysis, more accurate counting can also enable more accurate linking of tREs to their target genes with correlated counts, a growing focus of research ([2, 42, 43]).

The leading-edge leverages the consistency of the expression-based ranks to identify the set of tREs supported by both changes in transcription (the ranking) and co-localization of a responsive transcription factor (a supporting motif). We developed two approaches to identifying the leading-edge, i.e. the inflection point in motif to differential transcription enrichment. We found that the Plateaued Enrichment method provided the best calls of tREs with transcription changes specific to a TF. Conceptually, the leading-edge can also be used to assign responsibility for the observed expression change to a specific motif based on the motif’s co-localization. Yet these TF responsibility assignments are not necessarily exclusive, as each motif is analyzed independently of other motifs. Furthermore, while the leading-edge is calculated per motif, we show that the approach can also be applied to complex perturbations with more than one TF responding. Our reliance on motif instances, however, means that our assessment is limited by the accuracy and specificity of a TF’s motif. For example, we were unable to decipher between the activity of AhRR and AhR due to their essentially identical motifs.

The second method of identifying the leading-edge, the Match Background approach, proved highly informative on interpreting larger-scale enrichment trends that improve our ability to identify false positives in TF enrichments. We showed that metrics based on the leading-edge can distinguish between GC-corrected significant calls that show no clear enrichment and those with clear biological support from multiomics data or downstream validation. Indeed, with the leading-edge, subtle but biologically meaningful enrichment such as AhR within the woodsmoke data can now be detected without inflating the false positive rate[35]. Importantly, some TFs can function as both an activator and a repressor, dispersing their signals to both ends of the ranked list. Although the quantile plots provide a quick way to identify these TFs (high change at both poles), the TFEA-LE framework does not currently consider simultaneous enrichment in both directions. However, the leading-edge metrics do provide an intuitive interpretation of enrichment results, allowing users to consider a number of alternative enrichment cutoffs and their tradeoffs. Therefore, the leading-edge can be essential not only for improving the call of responsive tREs, but has largely improved TFEA calling of responsive transcription factors as well.

The leading-edge seems most advantageous when dealing with technically challenging multiomics data. In particular, TFEA-LE was robust to low overall transcription signal and outlier samples, cases that are problematic for classical statistical approaches. Finally, while our ranking is based entirely on expression data, any relevant data could be used for the ranking or as orthogol data, including ChIP-seq data or ATAC signal. As peak-based assays like ATAC-seq are particularly prone to technical factors that leads classic statistic tools to perform poorly [44], TFEA-LE holds great potential to improve analysis on these datasets. In this manner, TFEA-LE is relatively general and could be used in numerous data combinations to provide greater confidence in associating SNPs overlapping motifs to their functional impact.

Ultimately, this work has provided multiple, open-source tools within a novel pipeline for improved characterization of changes in expression at lowly transcribed regulatory elements. Our applications simultaneously improve identification of significantly changing tREs, linking tREs to their upstream regulators, defining coordinates of tREs, and providing further confirmation of transcription factors responsible for changing transcription. Together, this work clarifies key weaknesses in current regulatory element focused analyses, particularly with nascent RNA-sequencing, and provides several methods to advance their usage.

## 4 Code Availability

All code is available on Github and zenodo at the links provided below.

1. Modified LIET: https://github.com/Dowell-Lab/LIET/tree/LIET_EMGtoo,
2. Nextflow pipeline that can allow Mu Counts or fixed-window counting: https://github.com/Dowell-Lab/Bidir_Counting_Analysis,
3. Updated TFEA (with leading-edge and related metrics): https://github.com/Dowell-Lab/TFEA/tree/Lead_edge,
4. Source code for this work: https://github.com/Dowell-Lab/Improving_tRE_Analysis_Paper

## 5 Data Availability

All sequencing data for TP53 based analysis can be found in the NCBI Gene Expression Omnibus (GEO) under accession number GSE86222 (HCT116, SJSA, MCF7). Sequencing data for GR based analysis can be found under accession numbers GSE125623 (ChIP) and GSE124916 (GRO-seq). All SRRs used with annotations can be found in Supplemental Table 1.

## 6 Competing interests

Dr. Dowell is a founder of Arpeggio Biosciences. Drs Dowell and Allen are on a patent for inferring TF activity from eRNA activity (PCT/US2018/0182330). The remaining authors declare no competing interests.

## 7 Author contributions statement

M.A.A. and R.D.D. conceived of the original project. H.A.T. and R.D.D. designed the algorithms for Mu Counts and the edited format of LIET. R.D.D. conceptualized the Leading Edge and H.A.T. designed the final algorithms of it used in this work. H.A.T., R.D.D., and M.A.A. conceptualized how to evaluate the algorithms. H.A.T. implemented the algorithms, tested them, and analyzed data. J.T.S. confirmed accuracy of code and changes to LIET and provided general feedback. H.A.T. and R.D.D. wrote the manuscript which was then reviewed by all authors.

## 8 Acknowledgments

The authors first thank the anonymous reviewers for their valuable suggestions and editor for ensuring a smooth reviewing process. We also thank the alumni/current members of the Dowell lab, particularly Drs. Samuel Hunter and Rutendo Siguake, for their feedback on interpreting some early results.

## 9 Funding information

This work is supported in part by funds from the National Institutes of Health grant awards HL109557 and GM125871. The NSF NRT Integrated Data Science Fellowship (award 2022138) from the Biofrontiers Institute and the Curci Scholarship from the Shurl and Kay Curci Foundation (both to H.A.T.) also enabled some of this work.

## References

[1] Yao, L., Liang, J., Ozer, A., Leung, A.K.-Y., Lis, J.T., Yu, H.: A comparison of experimental assays and analytical methods for genome-wide identification of active enhancers. Nature Biotechnology, 1–10 (2022)

[2] Sigauke, R.F., Sanford, L., Maas, Z.L., Jones, T., Stanley, J.T., Townsend, H.A., Allen, M.A., Dowell, R.D.: Atlas of nascent RNA transcripts reveals enhancer to gene linkages. BMC Genomics in press (2025)

[3] Andersson, R., Refsing Andersen, P., Valen, E., Core, L.J., Bornholdt, J., Boyd, M., Heick Jensen, T., Sandelin, A.: Nuclear stability and transcriptional directionality separate functionally distinct RNA species. Nat Commun 5 (2014)

[4] Kundaje, A., Meuleman, W., Ernst, J., Bilenky, M., Yen, A., Heravi-Moussavi, A., Kheradpour, P., Zhang, Z., Wang, J., Ziller, M.J., Amin, V., Whitaker, J.W., Schultz, M.D., Ward, L.D., Sarkar, A., Quon, G., Sandstrom, R.S., Eaton, M.L., Wu, Y.-C., Pfenning, A.R., Wang, X., Claussnitzer, M., Yaping Liu, Coarfa, C., Alan Harris, R., Shoresh, N., Epstein, C.B., Gjoneska, E., Leung, D., Xie, W., David Hawkins, R., Lister, R., Hong, C., Gascard, P., Mungall, A.J., Moore, R., Chuah, E., Tam, A., Canfield, T.K., Scott Hansen, R., Kaul, R., Sabo, P.J., Bansal, M.S., Carles, A., Dixon, J.R., Farh, K.-H., Feizi, S., Karlic, R., Kim, A.-R., Kulkarni, A., Li, D., Lowdon, R., Elliott, G., Mercer, T.R., Neph, S.J., Onuchic, V., Polak, P., Rajagopal, N., Ray, P., Sallari, R.C., Siebenthall, K.T., Sinnott-Armstrong, N.A., Stevens, M., Thurman, R.E., Wu, J., Zhang, B., Zhou, X., Beaudet, A.E., Boyer, L.A., De Jager, P.L., Farnham, P.J., Fisher, S.J., Haussler, D., Jones, S.J.M., Li, W., Marra, M.A., McManus, M.T., Sunyaev, S., Thomson, J.A., Tlsty, T.D., Tsai, L.-H., Wang, W., Waterland, R.A., Zhang, M.Q., Chadwick, L.H., Bernstein, B.E., Costello, J.F., Ecker, J.R., Hirst, M., Meissner, A., Milosavljevic, A., Ren, B., Stamatoyannopoulos, J.A., Wang, T., Kellis, M.: Integrative analysis of 111 reference human epigenomes 518(7539), 317–330 10.1038/nature14248. Publisher: Nature Publishing Group. Accessed 2024-11-19

[5] Hunter, S., Sigauke, R.F., Stanley, J.T., Allen, M.A., Dowell, R.D.: Protocol variations in run-on transcription dataset preparation produce detectable signatures in sequencing libraries. BMC genomics 23(1), 1–18 (2022)

[6] Love, M.I., Huber, W., Anders, S.: Moderated estimation of fold change and dispersion for rna-seq data with deseq2. Genome Biology 15(12), 550 (2014)

[7] Rubin, J.D., Stanley, J.T., Sigauke, R.F., Levandowski, C.B., Maas, Z.L., Westfall, J., Taatjes, D.J., Dowell, R.D.: Transcription factor enrichment analysis (TFEA): Quantifying the activity of hundreds of transcription factors from a single experiment. Nature Communications Biology (2021) 10.1038/s42003-021-02153-7

[8] Jones, T., Sigauke, R.F., Sanford, L., Taatjes, D.J., Allen, M.A., Dowell, R.D.: TF profiler: a transcription factor inference method that broadly measures transcription factor activity and identifies mechanistically distinct networks 26(1), 92 10.1186/s13059-025-03545-2. Accessed 2025-09-04

[9] Danko, C.G., Hyland, S.L., Core, L.J., Martins, A.L., Waters, C.T., Lee, H.W., Cheung, V.G., Kraus, W.L., Lis, J.T., Siepel, A.: Identification of active transcriptional regulatory elements from GRO-seq data. Nat Meth 12(5), 433–438 (2015)

[10] Lidschreiber, K., Jung, L.A., Emde, H., Dave, K., Taipale, J., Cramer, P., Lidschreiber, M.: Transcriptionally active enhancers in human cancer cells. Mol Syst Biol 17(1), 9873 (2021)

[11] Maurano, M.T., Humbert, R., Rynes, E., Thurman, R.E., Haugen, E., Wang, H., Reynolds, A.P., Sandstrom, R., Qu, H., Brody, J., Shafer, A., Neri, F., Lee, K., Kutyavin, T., Stehling-Sun, S., Johnson, A.K., Canfield, T.K., Giste, E., Diegel, M., Bates, D., Hansen, R.S., Neph, S., Sabo, P.J., Heimfeld, S., Raubitschek, A., Ziegler, S., Cotsapas, C., Sotoodehnia, N., Glass, I., Sunyaev, S.R., Kaul, R., Stamatoyannopoulos, J.A.: Systematic localization of common disease-associated variation in regulatory DNA. Science 337(6099), 1190–1195 (2012)

[12] Chen, X.-F., Guo, M.-R., Duan, Y.-Y., Jiang, F., Wu, H., Dong, S.-S., Zhou, X.-R., Thynn, H.N., Liu, C.-C., Zhang, L., Guo, Y., Yang, T.-L.: Multiomics dissection of molecular regulatory mechanisms underlying autoimmune-associated noncoding SNPs 5(17), 136477 10.1172/jci.insight.136477. Accessed 2023-10-31

[13] Zaugg, J.B., Sahlén, P., Andersson, R., Alberich-Jorda, M., Laat, W., Deplancke, B., Ferrer, J., Mandrup, S., Natoli, G., Plewczynski, D., Rada-Iglesias, A., Spicuglia, S.: Current challenges in understanding the role of enhancers in disease 29(12), 1148–1158 10.1038/s41594-022-00896-3. Number: 12 Publisher: Nature Publishing Group. Accessed 2023-11-16

[14] Gally, F., Sasse, S.K., Kurche, J.S., Gruca, M.A., Cardwell, J.H., Okamoto, T., Chu, H.W., Hou, X., Poirion, O.B., Buchanan, J., Preissl, S., Ren, B., Colgan, S.P., Dowell, R.D., Yang, I.V., Schwartz, D.A., Gerber, A.N.: The muc5b-associated variant rs35705950 resides within an enhancer subject to lineage- and disease-dependent epigenetic remodeling. JCI Insight 6(2) (2021) 10.1172/jci.insight.144294

[15] Sasse, S.K., Dahlin, A., Sanford, L., Gruca, M.A., Gupta, A., Gally, F., Wu, A.C., Iribarren, C., Dowell, R.D., Weiss, S.T., Gerber, A.N.: Enhancer RNA transcription pinpoints functional genetic variants linked to asthma 16(1), 2750 10.1038/s41467-025-57693-x. Publisher: Nature Publishing Group. Accessed 2025-04-17

[16] Smith, G.D., Ching, W.H., Cornejo-Páramo, P., Wong, E.S.: Decoding enhancer complexity with machine learning and high-throughput discovery 24(1), 116 10.1186/s13059-023-02955-4. Accessed 2025-08-25

[17] Allen, M.A., Mellert, H., Dengler, V., Andryzik, Z., Guarnieri, A., Freeman, J.A., Luo, X., Kraus, W.L., Dowell, R.D., Espinosa, J.M.: Global analysis of p53-regulated transcription identifies its direct targets and unexpected regulatory mechanisms. eLife 3, 02200 (2014)

[18] Andrysik, Z., Galbraith, M.D., Guarnieri, A.L., Zaccara, S., Sullivan, K.D., Pandey, A., MacBeth, M., Inga, A., Espinosa, J.M.: Identification of a core TP53 transcriptional program with highly distributed tumor suppressive activity. Genome Research (2017) 10.1101/gr.220533.117

[19] Robinson, M.D., McCarthy, D.J., Smyth, G.K.: edgeR: a bioconductor package for differential expression analysis of digital gene expression data. Bioinformatics 26(1), 139–140 (2010) 10.1093/bioinformatics/btp616. Accessed 2023-11-22

[20] Law, C.W., Chen, Y., Shi, W., Smyth, G.K.: voom: precision weights unlock linear model analysis tools for RNA-seq read counts. Genome Biology 15(2), 29 (2014) 10.1186/gb-2014-15-2-r29. Accessed 2023-11-22

[21] Li, Y., Ge, X., Peng, F., Li, W., Li, J.J.: Exaggerated false positives by popular differential expression methods when analyzing human population samples 23(1), 79 10.1186/s13059-022-02648-4. Accessed 2025-08-28

[22] Buratin, A., Bortoluzzi, S., Gaffo, E.: Systematic benchmarking of statistical methods to assess differential expression of circular RNAs 24(1), 612 10.1093/bib/bbac612. Accessed 2025-08-28

[23] Azofeifa, J.G., Dowell, R.D.: A generative model for the behavior of RNA polymerase. Bioinformatics 33(2), 227–234 (2017)

[24] Wissink, E.M., Vihervaara, A., Tippens, N.D., Lis, J.T.: Nascent RNA analyses: tracking transcription and its regulation. Nature Reviews Genetics 20(12), 705– 723 (2019)

[25] Yao, L., Liang, J., Ozer, A., Leung, A.K.-Y., Lis, J.T., Yu, H.: A comparison of experimental assays and analytical methods for genome-wide identification of active enhancers 40(7), 1056–1065 10.1038/s41587-022-01211-7. Number: 7 Publisher: Nature Publishing Group. Accessed 2023-11-16

[26] Wang, Z., Chu, T., Choate, L.A., Danko, C.G.: Identification of regulatory elements from nascent transcription using dreg. Genome Research 29(2), 293–303 (2019) 10.1101/gr.238279.118

[27] Heinz, S., Benner, C., Spann, N., Bertolino, E., Lin, Y.C., Laslo, P., Cheng, J.X., Murre, C., Singh, H., Glass, C.K.: Simple combinations of lineage-determining transcription factors prime cis-regulatory elements required for macrophage and b cell identities. Molecular Cell 38(4), 576–589 (2010) 10.1016/j.molcel.2010.05.004. Accessed 2024-12-05

[28] Drexler, H.L., Choquet, K., Churchman, L.S.: Splicing kinetics and coordination revealed by direct nascent RNA sequencing through nanopores. Molecular Cell 77(5), 985–9988 (2020)

[29] Drexler, H.L., Choquet, K., Merens, H.E., Tang, P.S., Simpson, J.T., Churchman, L.S.: Revealing nascent RNA processing dynamics with nano-COP. Nature Protocols 16(3), 1343–1375 (2021) 10.1038/s41596-020-00469-y. Number: 3 Publisher: Nature Publishing Group. Accessed 2023-11-02

[30] Reimer, K.A., Mimoso, C.A., Adelman, K., Neugebauer, K.M.: Co-transcriptional splicing regulates 3′ end cleavage during mammalian erythropoiesis. Molecular Cell 81(5), 998–10127 (2021) 10.1016/j.molcel.2020.12.018. Accessed 2023-11-02

[31] Stanley, J.T., Barone, G.E.F., Townsend, H.A., Sigauke, R.F., Allen, M.A., Dowell, R.D.: LIET model: capturing the kinetics of RNA polymerase from loading to termination 53(7), 246 10.1093/nar/gkaf246. Accessed 2025-04-17

[32] Azofeifa, J.G., Allen, M.A., Hendrix, J.R., Read, T., Rubin, J.D., Dowell, R.D.: Enhancer RNA profiling predicts transcription factor activity. Genome Research (2018) 10.1101/gr.225755.117

[33] Subramanian, A., Tamayo, P., Mootha, V.K., Mukherjee, S., Ebert, B.L., Gillette, M.A., Paulovich, A., Pomeroy, S.L., Golub, T.R., Lander, E.S., Mesirov, J.P.: Gene set enrichment analysis: A knowledge-based approach for interpreting genome-wide expression profiles 102(43), 15545–15550 10.1073/pnas.0506580102. Publisher: Proceedings of the National Academy of Sciences. Accessed 2024-08-05

[34] Sasse, S.K., Dahlin, A., Sanford, L., Gruca, M.A., Gupta, A., Gally, F., Wu, A.C., Iribarren, C., Dowell, R.D., Weiss, S.T., Gerber, A.N.: Glucocorticoid-regulated bidirectional enhancer RNA transcription pinpoints functional genetic variants linked to asthma. medRxiv. Pages: 2022.11.10.22281906. 10.1101/2022.11.10.22281906. https://www.medrxiv.org/content/10.1101/2022.11.10.22281906v2 Accessed 2023-11-16

[35] Gupta, A., Sasse, S.K., Gruca, M.A., Sanford, L., Dowell, R.D., Gerber, A.N.: Deconvolution of multiplexed transcriptional responses to wood smoke particles defines rapid aryl hydrocarbon receptor signaling dynamics. J Biol Chem 297(4), 101147 (2021)

[36] Cardiello, J.F., Sanchez, G.J., Allen, M.A., Dowell, R.D.: Lessons from eRNAs: understanding transcriptional regulation through the lens of nascent RNAs. Transcription 11(1), 3–18 (2020)

[37] Sartorelli, V., Lauberth, S.M.: Enhancer RNAs are an important regulatory layer of the epigenome. Nat Struct Mol Biol 27(6), 521–528 (2020)

[38] Mousavi, K., Zare, H., Dell’Orso, S., Grontved, L., Gutierrez-Cruz, G., Derfoul, A., Hager, G.L., Sartorelli, V.: eRNAs promote transcription by establishing chromatin accessibility at defined genomic loci 51(5), 606–617 10.1016/j.molcel.2013.07.022. Publisher: Elsevier. Accessed 2023-11-30

[39] Hou, T.Y., Kraus, W.L.: Analysis of estrogen-regulated enhancer RNAs identifies a functional motif required for enhancer assembly and gene expression 39(11) 10.1016/j.celrep.2022.110944. Publisher: Elsevier. Accessed 2024-01-21

[40] Schaukowitch, K., Joo, J.-Y., Liu, X., Watts, J.K., Martinez, C., Kim, T.-K.: Enhancer RNA facilitates NELF release from immediate early genes 56(1), 29–42 10.1016/j.molcel.2014.08.023. Accessed 2023-11-16

[41] Pnueli, L., Rudnizky, S., Yosefzon, Y., Melamed, P.: RNA transcribed from a distal enhancer is required for activating the chromatin at the promoter of the gonadotropin α-subunit gene 112(14), 4369–4374 10.1073/pnas.1414841112. Accessed 2023-11-27

[42] Carullo, N.V.N., Phillips Iii, R.A., Simon, R.C., Soto, S.A.R., Hinds, J.E., Salisbury, A.J., Revanna, J.S., Bunner, K.D., Ianov, L., Sultan, F.A., Savell, K.E., Gersbach, C.A., Day, J.J.: Enhancer RNAs predict enhancer-gene regulatory links and are critical for enhancer function in neuronal systems. Nucleic Acids Res 48(17), 9550–9570 (2020)

[43] Shinkai, N., Asada, K., Machino, H., Takasawa, K., Takahashi, S., Kouno, N., Komatsu, M., Hamamoto, R., Kaneko, S.: SEgene identifies links between super enhancers and gene expression across cell types 11(1), 49 10.1038/s41540-025-00533-x. Publisher: Nature Publishing Group. Accessed 2025-08-15

[44] Gontarz, P., Fu, S., Xing, X., Liu, S., Miao, B., Bazylianska, V., Sharma, A., Madden, P., Cates, K., Yoo, A., Moszczynska, A., Wang, T., Zhang, B.: Comparison of differential accessibility analysis strategies for ATAC-seq data 10(1), 10150 10.1038/s41598-020-66998-4. Publisher: Nature Publishing Group. Accessed 2025-04-17

