## Supplemental Material for "Improving confidence of differential transcription calls in enhancers"

September 12, 2025


- [1] BioFrontiers Institute, University of Colorado, Boulder CO 80309 USA
- [2] Department of Molecular, Cellular and Developmental Biology, University of Colorado, Boulder CO 80309 USA
- [3] Department of Computer Science, University of Colorado, Boulder CO 80309 USA


#### Supplementary Notes

##### 0.1 Impact of classic statistical parameters

Although most parameter options for classic statistical tools had limited impact on recall or precision, the mean-dispersion estimation methods of DESeq2 had large implications on the tREs called significant according to F1 scores in all cell types (Supplemental Figure S2). Most consistently, the DESeq2 mean-based estimation method led to the worst recall and largely skewed p-value distributions; both markers of poor statistical capture of the data (Figure 1A, Supplemental Figure S27). While DESeq2 local-based dispersions increased recall and alleviated the skew of p-value calls, EdgeR and Limma had the most consistent p-value distributions without skew (Supplemental Figure S27).

### Supplementary Methods

### 1 Methods

All code for analyses are available at (https://github.com/Dowell-Lab/Improving_tRE_Analysis_Paper/) which will subsequently be referred to as **github:**.

#### 1.1 PRO-seq Analysis

##### 1.1.1 Trimming, Mapping, and Quality Control

Samples were mapped to the hg38 genome and NCBI RefSeq annotations were used (hg38 release GCF 000001405.40-RS 2023 03). All samples were trimmed and mapped using an in-house NextFlow pipeline (https://github.com/Dowell-Lab/Nascent-Flow), run with NextFlow (version 20.07.1). Briefly, fastq files were trimmed for adapter sequences and low quality bases using BBMap (version 38.05) and aligned to reference genomes with HISAT2 (version 2.1.0). Downstream mapped read files (CRAM files and IGV-compatible TDF files) were generated with Samtools (version 1.8), Bedtools (version 2.28.0), and IGVtools (version 2.14.1). Samples were then assessed for quality using metrics from the following software packages: FastQC (version 0.11.8), HISAT2 (version 2.1.0), Preseq (version 2.0.3), RSeQC (version 3.0.0), and BBMap (version 38.05).

##### 1.1.2 Identifying bidirectional transcripts

This approach was used for all datasets (separately for each dataset defined in Supplemental Table 1) unless otherwise noted. Regions of bidirectional nascent run-on transcription were identified using Tfit and dREG. For both, we removed multimapped reads and reads with low mapping quality score with the following code (all caps indicate a bash variable is being used):

```
samtools view -@ 16 -h -q 1 |${SRR}.bam | grep -P
```

where ${SRR} refers to the prefix of the bam file (usually the SRR key). For Tfit, we used the in-house NextFlow pipeline (https://github.com/Dowell-Lab/Bidirectional-Flow). Final analyses used $3'$ bedgraphs that were generated with the -3 flag of bedtools coverage. Briefly, Tfit was run in a two step process, first with the template

matching module to identify sites of bidirectional transcription, then these regions were used for input to fit the precise RNA polymerase behavior. For dREG, we followed the recommended pipeline (per https://github.com/Danko-Lab/dREG) and generated BigWig input files by converting the filtered BAM files using bedtools bamToBed to BED files, which where then converted to bedGraph format with bedtools genomecov, and finally BigWig files from bedGraphToBigWig.
Final consensus regions of bidirectional transcription were identified using *muMerge* version 1.1.0 (https://pypi.org/project/mumerge/). *muMerge* probabilisticly determines the most likely midpoint of transcriptional initiation for the bidirectional RNAs (mu or $\mu$). Briefly, Tfit and dREG bidirectional calls were first mumerged separately across all replicates. For p53, these regions were then merged again with each celltype noted as "conditions." For the GR and TNF cells, these regions were merged again with each paper being noted as "conditions." As done previously[15], the dREG and Tfit *muMerge* files were then combined such that calls above 2.5kb were removed and Tfit calls were used for any regions overlapping by at least 40% (relevant code found in (https://github.com/Dowell-Lab/Bidir_Counting_Analysis).

##### 1.1.3 Counting reads over regions

The counting pipelines used in this work can be found and run with a nextflow pipeline at (https://github.com/Dowell-Lab/Bidir_Counting_Analysis). Briefly, a bed file of consensus regions (bidirectionals) must be provided where the midpoints of regions are considered the centers of bidirectionals (e.g. from *muMerge*). Users define a fixed window from the midpoints of regions over which they want to consider tREs or gene TSS bidirectionals (e.g. 500 means total 1kb region). Gene bodies are counted over to exclude the gene TSS bidirectionals from gene counts as described previously [15]. The pipeline addresses overlapping transcription from nearby bidirectionals via the algorithm Mu_Counts. Briefly, bidirectional transcripts are counted strand-specifically where the transcript regions are from the center of the bidirectional (mu ($\mu$)) to which ever is shorter: the fixed window length or the $\mu$ of the next closest bidirectional. A full visualization of the pipeline can be found at Supplemental Figure S8. Counting over the fixed window length alone is also done so users have access to both. In both cases, to address overlapping transcription from genes, first, bidirectionals overlapping genes with at least a user-defined percentage of the gene-body region transcribed (according to bedtools coverage, default 70%) on both strands are removed. Counts for bidirectionals with transcribed genes overlapping one strand are replaced with the stranded counts from the strand without overlap multipled by 2. We then ensure that only the regions within genes predicted to not have overlapping transcription from bidirectionals are counted over. The longest isoforms of genes are used for final counts where the regions of bidirectionals with counts more than a user defined limit are removed from the region considered for that gene for counting.

#### 1.2 Differential Expression Benchmarking

The following packages were used with R version 4.4.0 (2024-04-24) on platform x86_64-apple-darwin20: DESeq2_1.44.0, edgeR_4.2.1, limma_3.60.4.

##### 1.2.1 Simulation based

Relevant code and figures for this section can be found at (**github:**/Simul_Bench_DE)

**Refraction Analysis** Two samples with $> 100M$ non-duplicated reads were used to assess the impact of tRE vs gene TSS counts with decreasing depths: SRZ1554311 and SRR1145801 ([6, 17]). SRZ1554311 is a combination of technical replicates with the curation detailed in [15]. tREs were identified from Tfit and *muMerge* was run on the two samples to ensure no overlapping bidirectionals. Full counting fixed windows of 600bp and 1kb were used. Bams with uniquely mapped and non-duplicate reads were subsampled using the -bs flag of samtools view with unique seeds for each subsample to ensure random variability.

**Dispersion Visualization** Estimated mean-dispersion trends and coefficients were calculated with DESeq2 default settings using the functions DESeq2DataSetFromMatrix, estimateSizeFactors, estimateDispersions, and nbinomWaldTest. This was performed on all three celltypes with Nutlin-3a/DMSO data (MCF7, HCT116, SJSA) and PRO-seq samples from HeLa cells perturbed with either dox-inducible shRNA Ints11 or dox-inducible shRNA control (latter had read depths above 70M and quality control scores of 1 according to DBNascent) ([5, 3, 2, 15]. Log fold changes between conditions (shRNA control vs

Ints11 and DMSO vs Nutlin) or biological replicates were visualized for gene TSS bidirectionals, gene bodies, and tREs. Counts were collected from genes as described above, while gene TSS bidirectionals and tREs used a 600bp, unstranded, fixed window. Counts for tREs overlapping a transcribed gene (>30 summed counts from samples) on both strands were removed, and if on only one of the strands were counted by multiplying the non-convoluted strand counts by 2. Any features with less than 21 counts between all samples within the same cell type were removed before analysis.

**Power Analysis** R package powsimR version 1.2.4 was installed from their github repository (https://github.com/bvieth/powsimR) according to their instructions. The HeLa cell samples and counts from Dispersion Visualization section were used for power analysis ([5]). Features were filtered at 16 rather than 21 counts. Parameters of the data were estimated by powsimR using a TMM normalization (estimateParam function). Differential transcription data was simulated with 25 simulations, 5% of features being differentially transcribed, and six different number of replicates (2, 3, 5, 10, 15, 20). The following normalization and differential expression software were used: Median Ratio Normalization and DESeq2, TMM and EdgeR-LRT (likelihood ratio test), TMM and EdgeR-QL (quasi-likelihood test), TMM and Limma-Trend, TMM and Limma-Voom. All simulations produced very consistent results so only DESeq2 with Median-Ratio normalization is shown.

##### 1.2.2 p53 Differential Expression Benchmarking

Relevant code and figures for this section can be found at (**github:**/Bench_DE) in subdirectories Truth_Sets and Before_LE.

**Tested Methods** The following combinations (total) were used to identify differentially transcribed regions: First, as a baseline, transcribed bidirectionals ($>$ 20 counts within cell type) with positive log fold changes were randomly assigned to “significant” or “not significant” with equal probability. For classic tools, the most common three tools were considered according to their relevant documentation. Details for these methodologies can be found in the documentation and papers corresponding to the appropriate packages [11, 14, 10]. DESeq2 was performed with local, mean, or parametric dispersion methods, Wald or Likelihood Ratio significant tests, and Ratio, positive counts, and iterative normalization methods.

EdgeR with Locfit, Movingave, LOESS, and Locfit.mixed dispersion methods (robust or not), Likelihood Ratio, quasi-likelihood, and quasi-likelihood-robust significance tests. Limma with voom or trend dispersion methods and eBayes or eBayes-robust significance tests. Both Limma and EdgeR used trimmed mean of M-values (TMM), TMM with singleton pairing (TMMwsp), upper-quartile, or relative log expression (RLE) for normalization methods. Finally, to further consider the impact of normalization on results, we considered virtual spike-in normalization factors [12]. Adjusted p-value cutoffs of $1e-70, 1e-50, 1e-30, 1e-20, 1e-18, 1e-16, 1e-14, 1e-12, 1e-10, 1e-8, 1e-6, 1e-4, 1e-2, 0.05, 0.1, 0.3, 0.5, 0.7, 0.9, 0.95$, and $0.99$ were used.

**Defining the Truth Sets** The expected "True Positives" were transcribed bidirectionals split into 4 sections for each cell type: BOTH (overlapping the appropriate cell type-specific ChIP peaks as called in their original, respective publications and having their bidirectional centers ($\mu$s) be within 500bp of the appropriate TF motif), EITHER (motif or CHIP qualifications), CHIP, and MOTIF. ChIP Peaks originally published in hg19 coordinates were converted to hg38 coordinates using UCSC Liftover (https://genome.ucsc.edu/cgi-bin/hgLiftOver) as available Spring 2024. To assess potential bias from motif prediction for p53, HOCOMOCOv12 motif P53.H12CORE.0.P.B was used with FIMO p-value significance cutoff of either p-value $1e-6$ or $1e-5$. The expected "True Negatives" were identified for each cell type as transcribed bidirectionals (total counts $> 20$ within cell type) not within 10kb of a ChIP peak, with below 11 standardized ($counts_{experimental} - counts_{control}$) ChIP peak reads, and with centers ($\mu$s) at least 8kb away from both p53 HCOCOMOCOv12 motifs (P53.H12CORE.0.P.B and P53.H12CORE.1.S.C) (FIMO p-value $1e-5$). The jupyter notebooks for getting these Truthsets can be found at P53_Classic_Bench_DE/Assess_TPs_H12.ipynb and Assess_TNs_H12.ipynb. Since ChIP peaks and motifs do not necessarily confer transcription[3], we also considered the three cell types as replicates to allow six replicates per condition rather than two. We considered the same tested differential expression methods explained above and had two groups to use as additional "True positive" sets: "Combined Union" and "Combined Intersect." The former are tREs called significant by any tested methods (N=1640). The latter are tREs called significant all tested methods except necessarily TMM_eBayes_Trend and TMM_eBayes-robust_Trend (for final

Combined Intersect N=399). These latter two methods were not required since they led to only 38 tREs being called across all methods.
Precision was calculated as TP/(TP+FP) where FP was all regions called significant that were considered a True Negative defined as above. Recall was calculated as the number of True Positives called by the tool divided by the number of True Positives defined above. Area under precision-recall curves usually provides a more robust evaluation of classification methods by considering all significance cutoffs. The differing ranges of precision and recall across platforms, however, prevented fair evaluation with this metric. Specifically, DESeq2 showed far lower maximum recall values compared to EdgeR and Limma for MCF7 and SJSA, even when a p-adjusted cutoff of 0.99 was included (Supplemental Figure S26). Adjusted pvalue cutoffs of $10^{-70}$, $10^{-50}$, $10^{-30}$, $10^{-20}$, $10^{-18}$, $10^{-16}$, $10^{-14}$, $10^{-12}$, $10^{-10}$, $10^{-8}$, $10^{-6}$, $10^{-4}$, 0.01, 0.05, 0.1, 0.3, 0.5, 0.7, 0.9, 0.95, and 0.99 were considered.

#### 1.3 Length based Benchmarking

Relevant code and figures for this section can be found at (**github:**/Length_Bench).

##### 1.3.1 Defining the Truth Length Sets

Long-read nascent run-on sequencing currently provides the best, high-throughput length estimation of enhancer-associated transcripts, but has few published experiments [8, 7, 13]. Therefore, we evaluated the ability of tools to predict the length of enhancer-associated transcripts (as defined by long-read nascent run-on sequencing) from short-read nascent run-on sequencing data. Five long-read nascent run-on sequencing fastq files (same control states) for K562 were downloaded from SRA with relevant information found in Supplemental Table 2 ([8]). Fastqs already had adapters and poly A/I tails removed according to Guppy as used by the original authors ([8]). Due to the low-depth of nascent long-read samples, and since individual reads rather than counts would be used, all fastqs were combined into a single fastq file before mapping. Following the code used by the original authors, reads were mapped to hg38 using minimap2 (primarily designed for error-prone long reads) with the following parameters (words in all caps refer to bash variables):

```
minimap2 -acx map-ont -t 16 -k14 \
--sam-hit-only ${FASTA} ${FASTQ} \
| samtools sort -o ${BAM}
```

where ${FASTA} points to the fasta file for the hg38 genome, ${FASTQ} refers to the long-read fastq file downloaded, and ${BAM} refers to the named bam file to serve as output.

As a truth set for comparison, 411 tRE associated transcripts with support from ENCODE Phase 3 (ENCFF464BRU) were manually annotated as transcriptionally isolated in both long-read and short-read data, having at least two long-reads (mapping quality $\geq 30$) supporting a clear transcript end position, and significant depth from previously published nascent run-on short read samples in K562 controls (SRA SRR4454567/8/9/70) [16, 1]. The most downstream end of long reads (minimum mapping quality of 30) within the annotated transcript region was used as the "true end" of the transcripts.

##### 1.3.2 Linking RNA calls across tools

Since a single tool might call multiple bidirectionals or transcripts within a region of interest, we identified the transcripts for each tool most appropriately aligning to the long-read supported transcripts with the following methods.

We used *muMerge* to get consensus bidirectional center calls for the relevant tools: dREG and Tfit. Briefly, coverage filtered (at least 9 counts per tRE predicted) output by dREG and Tfit were given to *muMerge* along with a metadata file grouping DMSO and heatshock samples together. For this work, a new flag was added to *muMerge* to allow the original positions of the regions and samples in which they're found to also be saved (–orig_names). The code run was

```
python ${SRC}/mumerge.py -i {METADATA} -o {OUT_PREFIX} \
--orig_names
```

where ${SRC} refers to the directory containing the cloned repository for *muMerge*, ${METADATA} is the file with the metadata information for the samples (e.g. replicates and conditions), and ${OUT_PREFIX} is the prefix for all output to be saved with. Users may get similarly coverage filtered regions and details on parameter layouts using the verbose branch of *muMerge*: https://github.com/Dowell-Lab/mumerge/tree/verbose.

Tfit has been shown to have the highest accuracy in calling the center of bidirectional transcription and using the $3'$ bedgraphs particularly improves the calls (this work) ([15, 18]). Therefore, the final centers of bidirectional transcription ($\mu$s) were based on Tfit $3'$ calls. Bedtools closest was run to find the distances between Tfit $3'$ calls and dREG and Tfit calls. A bidirectional was then linked to the original Tfit $3'$ bidirectional of interest if the $\mu$s (center of bidirectional) were within 500bp of one another. Homer calls were matched in two ways: 1) requiring a transcript's $5'$ end to be within 400bp of the Tfit $3'$ $\mu$, or 2) requiring a transcript to overlap region between Tfit $3'$ $\mu$ and downstream 100bp and in the case of multiples, choose the one where the $5'$ end was closest to $\mu$. The latter method includes transcripts that have poor calls for the transcript start but still capture the transcript. dREG and Tfit (full read) calls were matched to Tfit $3'$ calls according to the $\mu$s being within 500bp.

##### 1.3.3 Noise generation

To determine the regions over which to add noise, we used 1.5kb downstream of the true length annotation (according to long read) with a minimum of 3.5kb downstream of the bidirectional center ($\mu$) [padded regions]. If a strand of a bidirectional does not have any long reads, then 2kb is used. All regions were manually examined to ensure significant reads, short or long, were not omitted and nearby transcripts were not accidentally included. To assess the impact of noise, we added reads in random locations (sampling with replacement) across the padded regions. The number of reads added was different percentages of the number of original reads mapped to the same region (filtering for supplementary or secondary alignment): 20, 40, 60, 80, 100, 120, and 140. To mimic full reads, the random position was elongated to the $5'$ end by 74bp to reach the original 75bp read length. To mimic the $3'$ ends of reads in a bedgraph, the position alone was used. These reads (formatted as bedgraphs) were then merged with the original multi-mapped filtered bedgraphs using bedtools version 2.28.0.

##### 1.3.4 Homer

Homer v5.1 was run with multi-mapped filtered bams or bedgraphs (full reads) ([9]). First, tag directories were made using 'makeTagDirectory' and the '-keepAll' flag to ensure the same input was used across all tools. Then, nascent run-on sequencing de novo transcript identification was run with (bash variables in all caps)

```
findPeaks ${TAGDIR} -style groseq -o ${OUT_FILE} \
-minBodySize 150 -tssSize 50 -bodyFold 3 \
-endFold 5 -uniqumap hg38-50nt-uniqmap
```

where ${TAGDIR} refers to the TagDirectory from the filtered bams/bedgraphs, ${OUT_FILE} is the output path to save the peaks. Different parameter combinations like endFold 7, bodyFold 4, and tssFold 4 were also tested with the above showing the best results and therefore used for benchmarking. More detailed explanation of the Homer groseq algorithm can be found at http://homer.ucsd.edu/homer/, hereby called **homer:** at homer:/ngs/groseq/groseq.html where you can also download the hg38-50nt-uniqmap folder at homer:data/uniqmap/uniqmap.hg38.50nt.zip. The latest version available at the time of this work was October 26, 2018.

###### 1.3.5 dREG

dREG uses single-position based bigwigs. To convert the $3'$ bedgraphs to bigwigs, we used bedGraphToBigWig v4. We then ran dREG v1.0 through https://dreg.dnasequence.org/ using default parameters (R version 4.3.2 (2023-10-31)). The dREG lengths are directly based on dREG output.

###### 1.3.6 Tfit

For initial analysis, Tfit was run with both $3'$ bedgraphs or full read bedgraphs using Tfit_focus branch of the pipeline in https://github.com/Dowell-Lab/Bidirectional-Flow/. This nextflow pipeline has been optimized to run Tfit genome-wide. To allow unnecessary computational burden, for noise based calls, we had Tfit only search over the regions of study by feeding Tfit the padded regions used for noise production as preliminary regions. Tfit was run using the shell script available at https://github.com/Dowell-Lab/Bidirectional-Flow/blob/main/bin/tfit_model.sh (commit 89f2fd1, words in all caps are bash variables) :

```
${TFIT_DIR}/tfit_model.sh -t ${TFIT_PATH} \
-c ${TFIT_CONFIG} -b ${BG} -k ${PADDED_REGIONS} \
-p ${PREFIX} -n 32
```

where ${TFIT_DIR} refers to the path where the tfit_model.sh script is, ${TFIT_PATH} is the path to the Tfit software, ${TFIT_CONFIG} is the path to the config file for Tfit, ${BG} is the bedgraph used by Tfit, ${PADDED_REGIONS} is a bed file with the padded regions used for noise calls (considered preliminary regions over which to look for bidirectionals), and ${PREFIX} is the prefix used to name output. Details regarding the parameters can be found at https://github.com/Dowell-Lab/Bidirectional-Flow/. Tfit release version 1.2 (Repository version) was used and can be found at https://github.com/Dowell-Lab/Tfit/releases/tag/v1.2. Importantly, in this version, the source code incorrectly labels $\tau$ as $\lambda$. To avoid confusion, all cases where Tfit's $\lambda$ parameter is used, but is actually $\tau$, we label as $\tau$ here.

Tfit lengths were considered according to three options:

1. The original output length from Tfit (when going from $\mu$ to the edge of the region): $\tau + \sigma$,
2. $\tau + \sigma + \frac{\text{footprint}}{2}$, since the footprint is not originally considered in the length output of the region from Tfit, and
3. $\frac{\text{footprint}}{2} + |x - \mu|$ where $x$ is the 95th percentile of the EMG, numerically solved for given $\mu$, $\sigma$, $\tau$

   $$CDF(x \mid \mu, \sigma, \tau) = 0.95$$

   where $CDF(\cdot)$ is the cumulative distribution function of the EMG and $\tau$, $\mu$, $\sigma$ are parameters of the EMG.

##### 1.3.7 LIET

LIET v1.0.0 takes both a pad file to determine the complete regions over which to look and annotation file to use as priors in its Bayesian modeling. We used the same pads used for noise production (1.5kb downstream of the true length annotation (according to long read) with a minimum of 3.5kb downstream of the bidirectional center ($\mu$)). Input annotations for LIET were entered with the $5'$ location being the bidirectional center from *muMerge* ($\mu$) + Tfit footprint/2 as the start and 400bp downstream of this position as the $3'$ location, labeling all regions as positively stranded for simple tracking purposes. Exact LIET parameters and priors used are found in Supplementary Table 1. Originally, LIET was designed to run the full LIET model on the sense strand (positive according to the annotation) and an EMG model on the antisense strand. The software was edited to run either the EMG or LIET model on either strand. Similarly, percentiles of the probability distributions (with background removed) were calculated. Briefly, weights making up the total weight ($w$) were recalculated (to $w'$) after removing background ($w_b$) as described by Equation 1. The probability distribution functions of each strand using these recalculated weights were then calculated based on a domain of $(-10^6, 10^6)$. As defined in Equation 2, we compute the genomic positions corresponding to each target percentile ($q$) from these pdfs by identifying where the strand-specific cumulative distributions reach that value.

$$\text{Given } \mathbf{w} = [w_1, w_2, \ldots, w_{n-1}, w_b], \quad \text{define } \mathbf{w}' = \left[\frac{w_1}{\sum_{i=1}^{n-1} w_i}, \frac{w_2}{\sum_{i=1}^{n-1} w_i}, \ldots, \frac{w_{n-1}}{\sum_{i=1}^{n-1} w_i}\right] \tag{1}$$

where:

- w = original vector of weights for the LIET model,
- w′ = recalculated vector of weights for the LIET model.

$$\begin{aligned} \text{CDF}_p(x) &= \sum_{i=1}^{x} \text{PDF}_p(i) \\ \text{CDF}_n(x) &= \sum_{i=1}^{x} \text{PDF}_n(i) \\ \text{position}_p(q) &= \arg\min_x |\text{CDF}_p(x) - q| - 10^6 \\ \text{position}_n(q) &= \arg\min_x |\text{CDF}_n(x) - q| - 10^6 \end{aligned} \tag{2}$$

where:

- $\text{CDF}_p(x)$ = cumulative distribution for the positive ($p$) strand,
- $\text{CDF}_n(x)$ = cumulative distribution for the negative ($n$) strand,
- $\text{position}_p(q)$ = closest position to percentile $q$ on the positive strand (0-based),
- $\text{position}_n(q)$ = closest position to percentile $q$ on the negative strand (0-based).

##### 1.3.8 Calculating consensus lengths

To calculate consensus lengths based on replicates similar to *muMerge*, we considered multiple methods. First, we took the average. Next, we took a weighted average according to weights of coverage or in the case of LIET, the number of reads assigned outside of background:

$$\text{Weighted Average } 3' \text{ position} = \frac{\sum_{i=1}^{n} x_i \cdot c_i}{\sum_{i=1}^{n} c_i}$$

where $x_i$ is the $3'$ position of the transcript in sample $i$ and $c_i$ is either full coverage of the transcript according to feature counts or the weight of transcription $(1 - w_b)$ according to LIET in sample $i$.

#### 1.4 Length based p53 Differential Expression Benchmarking

Relevant code and figures for this section can be found at (**github:**/Bench_DE/Length_DE).

##### 1.4.1 Mu_Counts

This description follows what is shown in Supplemental Figure S8. The full pipeline (as a Nextflow pipeline) and additional descriptions are accessible at https://github.com/Dowell-Lab/Bidir_Counting_Analysis/tree/main. The first step involves filtering inputs. First, the tool takes in consensus regions for bidirectional transcripts where the midpoint is assumed to be the initiation point of bidirectional transcripts and the length is a confidence interval around it (like from *muMerge*). Therefore, we remove bidirectionals that are likely called due to technical noise (those with confidence intervals above 3.5kb) and filter bams/crams to remove multimapped reads. This module produces consensus files with the widths needed for future analyses and unique names (based on parameters). The next major step is identifying Gene TSS Bidirectionals; these show distinct transcriptional patterns from tREs and correspond to gene PROMPTs rather than enhancers. Bidirectionals coordinating to the transcription start site (TSS) of genes are identified by overlapping bidirectionals with a small window (parameter TSS_WIN - default 25bp) with 1kb regions around gene TSSs (parameter tss_1kb_file, also available for GR38.p14). Gene TSS bidirectionals are assigned so that a single gene isoform only has one TSS bidirectional, with the one whose midpoint is closest to the TSS used. Multiple gene isoforms are allowed to share the same TSS bidirectional if their TSSs are within 50bp of each other. The third step involves addressing overlapping transcription from both genes and other tREs. Nascent transcription of genes continues downstream of annotations and many tREs are found within introns. Therefore, gene bodies considered transcribed (parameter COUNT_LIMIT_GENES=70% isoform is covered with reads) along with the region 10kb downstream of said isoform are overlapped with the nonTSS bidirectionals (tREs). Any nonTSS bidirectional overlapping active gene transcription on both strands is removed since deconvolution cannot confidently occur. NonTSS bidirectionals overlapping gene transcription on one strand have counts replaced with those from the strand with no overlapping transcription, doubled. We then address overlapping transcription from bidirectionals with other bidirectionals using Mu_Counts. If the parameter COUNT_WIN for a tRE means the tRE is now overlapping with another tRE region, the neighborhing tREs will be counted so that the maximum distance of each RNA is the $\mu$ of the nearest neighboring tRE. RNAs for tREs are then counted separately according to the strand (e.g. counts on positive/negative strand for a bidirectional's positive/negative RNA are counted separately before being combined). Finally, we address gene counts that improperly contain counts from overlapping bidirectionals. Azofeifa et al[4] showed that gene counts can be largely disrupted by including responding bidirectionals. Therefore, the pipeline optionally removes the regions of bidirectionals with counts above parameter COUNT_LIMIT_BIDS.

**Grouping Test Sets** We hypothesized that length would have testable implications on two key groups of calls: Isolated (bidirectionals without possible convolution from nearby transcription) and "Overlapping" pairs ("True positive" and "True negative" bidirectionals that can influence each other's calls with convoluting transcription). Isolated bidirectionals were identified as those with no other tREs with 5.5kb of them, no overlap with genes, and at least 500bp upstream of a gene TSS and 10kb downstream of a gene annotated termination site (only considering genes that had total counts $> 200$). Overlapping pairs were identified as "True positives" and "True negatives", as defined below, with $\mu$s within 5kb of each other.

**Defining the Truth Sets** In order to not bias the truth sets to a specific tool, we took a slightly different approach to defining the truth sets for this length-based analysis compared to the analysis focusing on differential transcription alone. First, we wanted to limit how much the position of $mu$ potentially bias truth sets as Homer cannot consider $mu$. Therefore, we considered "True Positives" if a 50bp region around Tfit $3'$ $\mu$

overlapped a ChIP peak with a p53 motif within it. To ensure we had high enough statistical power when considering overlapping "True Positives" and "True Negatives," we used less stringent requirements for these truth sets when considering overlapping tREs. Expected "True Positives" were considered according to p53 ChIP peaks containing TP53 HOCOMOCOv12 motifs with a p-value cut off of 1e-5. Expected "True Negatives" were transcribed bidirectionals (total counts > 20) without motifs or ChIP peaks within 2kb rather than 10kb to allow consideration of "Overlapping" pairs within 5kb of each other.

#### 1.5 TFEA and Leading Edge Analysis

Transcription Factor Enrichment Analysis from https://github.com/Dowell-Lab/TFEA (version v1.1.1) was run with the ranked files (according to log fold change, then adjusted p-values) from the different parameter-tool combinations. The FIMO (from Meme v5.0.3) scanning background was set to uniform. Otherwise, default parameters were used.

##### 1.5.1 Leading Edge Methodologies

The final algorithm is integrated into the most updated version of TFEA (v2.0.1): https://github.com/Dowell-Lab/TFEA. Relevant code and figures from using the leading edge for identifying significantly changing tREs in p53 and GR datasets can be found at for this section can be found at (**github:**/Bench_DE) in subdirectories Get_LE and Compare_LE. Simply speaking, the leading edge is interpreted the position at which any elements with higher p-values (hence less statistically significant changes) are no longer considered as contributing to the transcription factor being enriched. Two methods to find the leading edge were used:
Let $E(t)$ represent the cumulative enrichment as a function of the ranked tREs $t$.

1. **Matched Background:** The first position (so leftmost if positive enrichment score and rightmost if negative) where the slope of the cumulative enrichment score curve is equal to or below that of the background enrichment line.

   Let $B(t)$ represent the background enrichment line (a straight line from $E(0)$ to $E(T)$).

$$\text{Slope of enrichment:} \quad E'(t)$$

$$\text{Slope of background:} \quad B'(t) = \frac{E(T) - E(0)}{T}$$

Define the matched background position $t^*$ as:

$$t^* = \begin{cases} \min\{t : E'(t) \leq B'(t)\} & \text{if } auc(E(T)) > 0 \text{ (positive enrichment)} \\ \max\{t : E'(t) \geq B'(t)\} & \text{if } auc(E(T)) < 0 \text{ (negative enrichment)} \end{cases}$$

2. **Plateaued Enrichment:** Captures the position at which the cumulative enrichment changes have stopped steadily changing due to enrichment. The second derivative of cumulative enrichment ($E''(t)$) shows two cases: a monotonic stabilization towards 0 or a non-monotonic function with up to six extrema (high oscillation) before stabilization. With non-monotonic cases, we use the first extrema as a conservative leading edge. With monotonic stabilizing cases (no extrema within the first 40% of tREs), we calculate the elbow of the curve as shown below.

$$t^* = \begin{cases} \arg\min_{t<0.4T} E'''(t) = 0 & \text{(Early Oscillation)} \\ \text{elbow}(E''(t)) & \text{(Monotonic stabilization)} \end{cases}$$

where $T$ is the total number of tREs and the elbow is computed as:

$$\text{elbow}(E''(t)) = \arg\max_t (\text{distance from line } \ell(t) \text{ from } E''(0) \text{ to } E''(T))$$

**Smoothing Method:** To reduce sensitivity to noise, multiple smoothed iterations of the cumulative enrichment curve $E(t)$ are generated using B-spline interpolation with degree $k = 5$. The final leading edge is determined by taking the median position across all spline fits. The initial smoothness parameter and subsequent smoothness sequences for the spline were optimized empirically by assessing about 100 case-scenarios: across several perturbations (P53, TNF/DEX, WSP, UPM, shRNAs) and cell types (HCT116, MCF7, SJSA, BEAS2B, different primary samples of small airway epithelial cells, ESC), bidirectional numbers ranging from 15,000 to 120,000, and transcription factors with motifs covering from 0-50% of tREs. Importantly, subsequent smoothness assessment (described

in next section) ensures that the algorithm is robust to this initial smoothness parameter.
Let:

- $N_{\text{motif}}$: number of tREs with motif calls
- $s_0$: initial smoothness value, defined by a tiered rule based on $N_{\text{motif}}$
- $s_i$: spline smoothness parameter for iteration i

**Initial smoothness $s_0$:**

$$s_0 = \begin{cases} 3 \times 10^{-12} & \text{if } N_{\text{motif}} > 30000 \\ 4 \times 10^{-12} & \text{if } N_{\text{motif}} > 20000 \\ 5 \times 10^{-12} & \text{if } N_{\text{motif}} > 10000 \\ 2 \times 10^{-11} & \text{if } N_{\text{motif}} > 8000 \\ 3 \times 10^{-11} & \text{if } N_{\text{motif}} > 7000 \\ 4 \times 10^{-11} & \text{if } N_{\text{motif}} > 5000 \\ 5 \times 10^{-11} & \text{if } N_{\text{motif}} > 4000 \\ 7 \times 10^{-11} & \text{if } N_{\text{motif}} > 3000 \\ 8 \times 10^{-11} & \text{if } N_{\text{motif}} > 2500 \\ 9 \times 10^{-11} & \text{if } N_{\text{motif}} > 2000 \\ 4 \times 10^{-11} & \text{if } N_{\text{motif}} > 1500 \\ 1 \times 10^{-10} & \text{if } N_{\text{motif}} > 1000 \\ 2 \times 10^{-10} & \text{otherwise} \end{cases}$$

**Smoothness sequence:**
Subsequent smoothness values are decreased to allow less smoothing:

$$s_{i+1} = s_i + 5 \times 10^{-(n_i - 1)}, \quad \text{where } n_i = \text{power of the current smoothness parameter}$$

This continues until the number of extrema for $E''(t)$ ($E'''(t) = 0$) exceeds a threshold (with a maximum of 20 iterations):

$$\text{Max } \left(E'''(t) = 0\right) = \begin{cases} 6 & \text{if } N_{\text{tRE}} > 60{,}000 \\ 4 & \text{otherwise} \end{cases}$$

Importantly, the ultimate leading edge is robust to these thresholds (4, 5, or 6), but we found that these separations ensured the greatest breadth of smoothness parameters considered while avoiding noise.

**In case of under-smoothed start:**
If the initial $s_0$ already exceeds the allowed number of extrema, then it is increased by $3 \times 10^{-(n_0)}$ until the constraint is satisfied:

$$s_0 = s_0 + 3 \times 10^{-(n_0)}, \quad \text{where } n_0 = \text{ power of the initial smoothness parameter}$$

##### 1.5.2 Other Updates to TFEA:

Relevant code and figures for this section can be found at (**github:**/Improving_FP_calls).

**TF-specific FIMO significance cutoffs:** To determine good default adjusted p-value cutoffs for FIMO (using Meme v5.0.3) for each motif, all 847,522 bidirectionals identified in [15] were scanned for motifs across their 3kb total regions (± 1.5kb from $\mu$) and using p-value cutoffs of $1e-4, 1e-5, 1e-6$, and $1e-7$. If a motif was called in between .5% (4,237) and 7% (59,326) bidirectionals based on one specified p-value cutoff, that p-value was considered the default cutoff for the TF to use. If less than .5% of bidirectionals had a motif at $1e-7, 1e-6, 1e-5$, or $1e-4$, the default p-value cutoff was changed to $1e-6, 1e-5, 1e-4$, and $1e-3$, respectively. If greater than 5% of bidirectionals had a motif at $1e-4, 1e-5, 1e-6$, and $1e-7$, the default p-value cutoff was changed to $1e-5, 1e-6, 1e-7$, and $1e-8$, respectively. If greater than 40% of bidirectionals had a motif at $1e-7$, the default p-value was changed to $1e-9$. Transcription factors with more than 20,000 tRE motifs for tREs found within the Nutlin-3a and DMSO dataset with $1e-6$ (ZN121, ZN135, ZN441, ZN560, ZN613, ZN770) were assigned best pvalues of $1e-10$. The code for this analysis can be found at Other_TFEA_Updates/Assess_pval_motifs.ipynb. TFEA was edited to take in a file with the desired p-value motifs with the option –fimo_thresh and the default values calculated above are provided as a file in the github repository of TFEA.

**Leading Edge metrics for significance calls:** False positives and true negatives were defined as the TFs with the highest enrichment scores that had no well-characterized linkage to the perturbation and were called significant or not according to the GC-corrected adjusted-pvalue ($< 0.01$), respectively. TFs with enrichment scores below 0.05 were not considered as they could be easily filtered out. Three metrics were calculated regarding the tREs with changes in enrichment that were higher than that from

background. Background slope was calculated as the maximum(cumulative enrichment score) - minimum(cumulative enrichment score) / Number of tREs. Frac_Background, as focused on in the remainder of the text, is the fraction of tREs with their slope higher than background. The fraction of tREs in the interquartile range of ranks with slopes higher than background (Frac_Back_Q13) and the difference between slopes (True - Background) (Diff_Back) were also calculated and showed comparable trends to Frac_Background.
To identify the quantile of ranked tREs where the cumulative enrichment score increased the fastest in magnitude, tREs were binned into 15 quantiles (around 3000 tREs per bin). The quantile with the maximum absolute value slope when using the median spline tested was returned as the Max_Quant. TFs with Max_Quant values in the middle of the ranked list (6,7,8,9,10) were considered False positives by the LE.

###### 1.5.3 Wood Smoke Particle LE and TFEA-LE Analysis:

Relevant code and figures for this section can be found at (**github:**/WSP).

**Matching ATAC-seq peaks and PRO-seq tREs:** To allow complete comparison between ATAC-seq and PRO-seq, bidirectionals were first called within each sequencing approach with Tfit before being mapped to each other. 80bp windowed PRO-seq bidirectionals (muMerged according to condition (BID)) were mapped to 1kb windowed ATAC-seq peaks (muMerged according to condition (ATAC)) with bedtools closest (words in all caps refer to variables):

```
bedtools closest -k 4 -d -D "ref" -a ${BID} -b ${ATAC} > ${OUT}
```

where ${BID} refers to the bed file of muMerged bidirectionals from PRO-seq and ${ATAC} refers to the muMerged ATAC-peak bed file. PRO-seq bidirectionals with $\mu$s within 2kb of an ATAC peak $\mu$ were kept with the closest corresponding ATAC peaks removed. 74% of PRO-seq bidirectionals (50,850) mapped to 60% (50,512) of ATAC-peaks. Otherwise, ATAC peaks and PRO bidirectionals were kept and noted as only occurring in one method. All regions were used for downstream analysis.

**Running TFEA:** To ensure the fairest comparison between ATAC-seq and PRO-seq, we only considered Non-GeneTSS bidirectionals/peaks as defined by Counting_Bid_Analysis. For TFEA and leading edge, rankings from EdgeR-TMM-QL were used but DESeq2 and other EdgeR results showed the same trends noted in this work. TFEA was run with and without assuming a uniform background for FIMO and led to no clear difference in results.

**Considering Leading Edge Metrics for TF Calls:** To ensure results weren't solely based on the number of motifs, TF calls were first filtered to have between 600 and 10000 tREs with the motif. To be considered a call supported by the leading edge metrics, the Match-Background leading edge had to be before the midpoint and after the Plateaued Enrichment leading edge, the Fraction of tREs above background had to be below 0.46 for PRO-seq and 0.51 for ATAC-seq, and the Max_Quant could not be 6,7,8,9,10 (interquartile range inclusive). TFs were then split into the following categories:

- **All**: GC-corrected Padj $< 0.001$, uncorrected Padj $< 0.001$, and meets LE requirements
- **GC_only**: GC-corrected Padj $< 0.001$, uncorrected Padj $\geq 0.001$, does not meet LE requirements
- **UNC_only**: GC-corrected Padj $\geq 0.001$, uncorrected Padj $< 0.001$, does not meet LE requirements
- **GC_LE**: GC-corrected Padj $< 0.001$, uncorrected Padj $\geq 0.001$, and meets LE requirements
- **UNC_LE**: GC-corrected Padj $\geq 0.001$, uncorrected Padj $< 0.001$, and meets LE requirements

The jupyter notebook with this analysis can be found at WSP/Plot_MB_curves.ipynb
The TF calls were assessed for directionality by comparing GC-corrected Enrichment scores. If the magnitude of scores were both $> 0.05$, they were assessed as being in the same or opposite directions ($\pm$) between ATAC-seq and PRO-seq of the same time points, or when using gene TSS bidirectionals (according to gene differential rankings) compared to tREs in the same condition.

#### Supplementary Figures

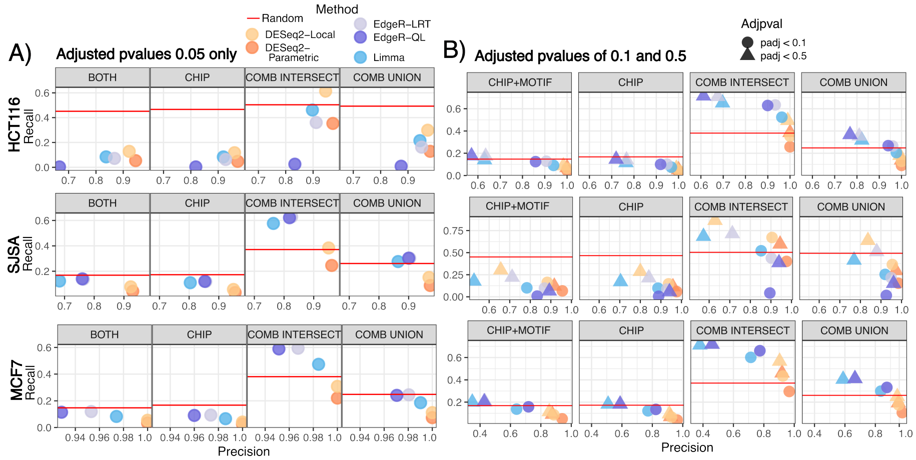


Figure S1: **High adjusted p-value cutoffs of 0.1 or 0.5 are required to reach recall of p53 truth sets enabled from random calling.** Recall and Precision for Nutlin-3A (p53) responding tREs when using five different classic statistical method combinations. True positives are based on p53 ChIP peaks (CHIP) or peaks with p53 motifs (BOTH) or calls achieved by all tools/parameter combinations (COMB INTERSECT) or any tool/parameter combination (COMB UNION) when considering cell types as replicates. False positives are based on calls without both motif and P53 ChIP peak. Red lines indicate recall from randomly assigning tREs with positive fold changes as a true call. A) Adjusted p-values $< 0.05$ are used. B) Adjusted p-value cutoffs of 0.1 or 0.5 are used.

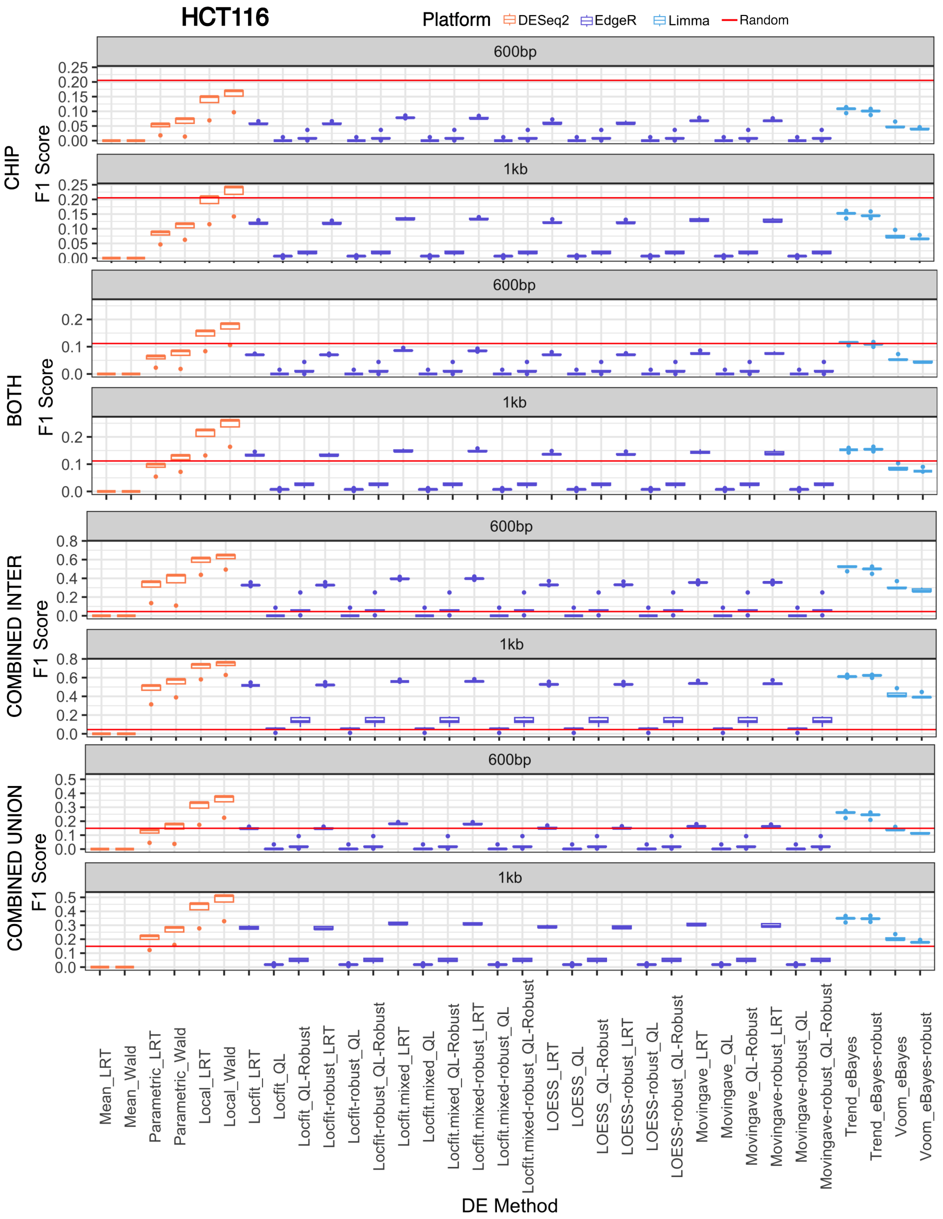


Figure S2: **Low F1 scores, specifically for the DESeq2-mean parameter combination, are consistent across varying truth sets across cell types.** A. HCT116 cells (Continued on the following page.)

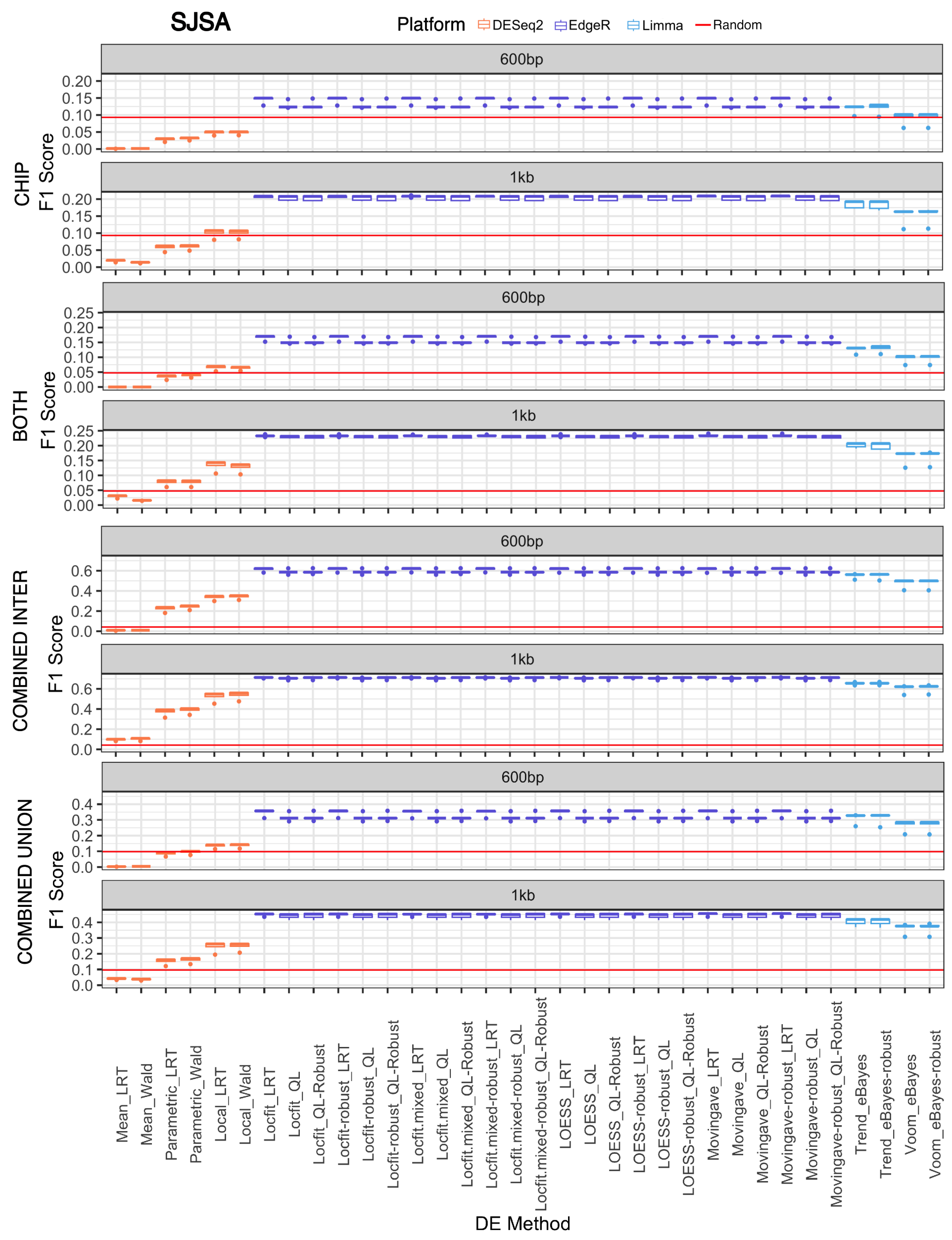


Figure S2: B. SJSA cells (Continued on the following page.)

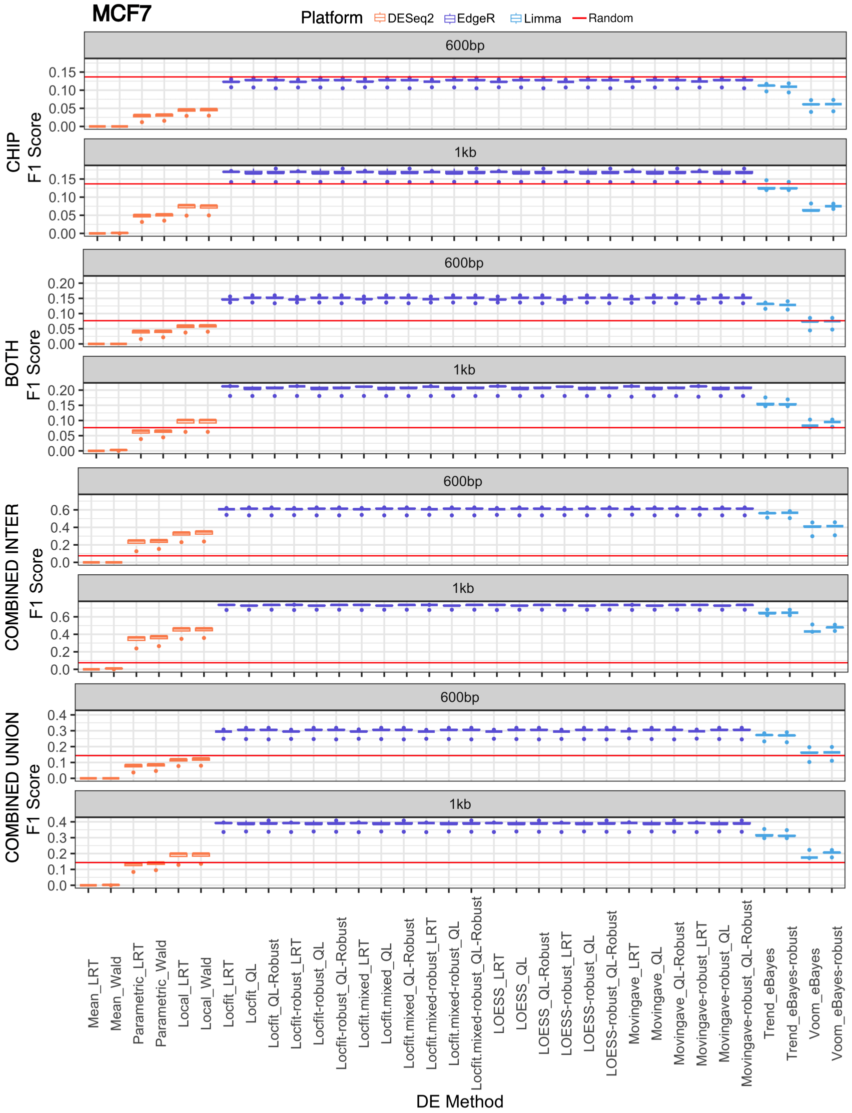


Figure S2: C. MCF7 cells. (A-C) F1 scores for all tested dispersion-significance test combinations are shown as boxplots colored according to the platform (DESeq2, EdgeR, or Limma). The red horizontal lines refer to the median F1 score calculated when randomly assigning features as significant five times. 600bp and 1kb refer to the window sizes used over which to count tREs. BOTH, CHIP, Combined Intersect, and Combined Union refer to the true positive sets used (details in Supplementary Methods).

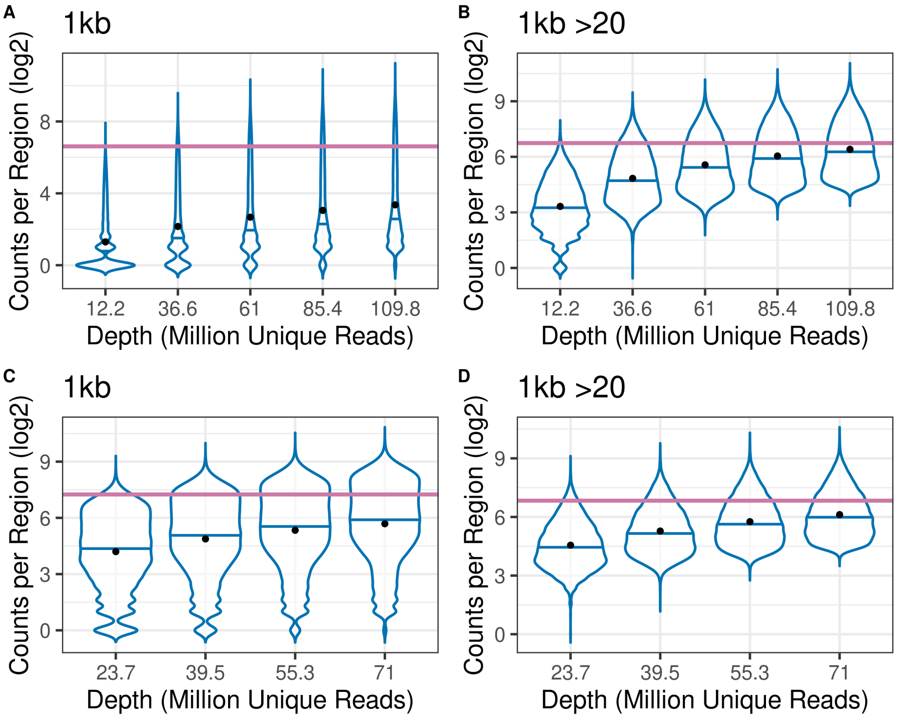


Figure S3: **About 100 million unique reads are needed for tREs to reach the same median counts obtained by gene TSS bidirectionals when only using about 40 million unique reads across two independent samples.**. Distribution of counts, including all tREs with counts above 0 (A+C) or 20 (B+D) at full depth visualized as violin plots. The median counts for gene TSS bidirectionals at 36.6 million (A+B) or 39.5 million (C+D) unique reads is shown as a pink line. Data in A+B is SRZ1554311 and Data in C+D is SRR1145801.

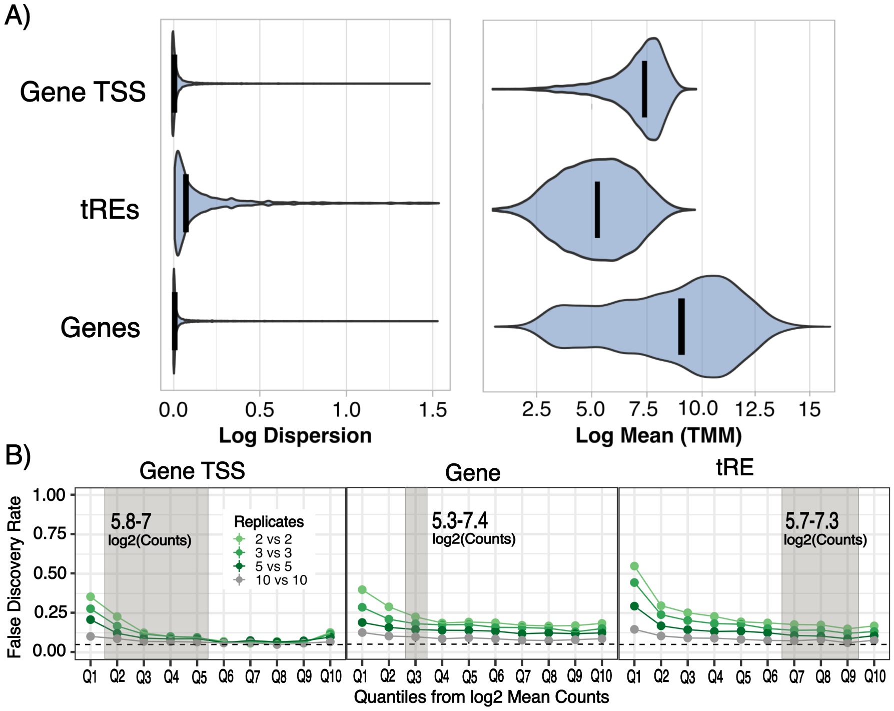


Figure S4: **Simulated data with mean-dispersion trends comparable to p53 data show that tREs require much higher number of replicates to achieve comparable false discovery rates as genes and gene TSS bidirectionals. A.** Distributions were estimated by powsimR based on real PRO-seq count data of the following features from two biological replicates (details in Supplementary Methods). **B.** False Discovery Rates of gene TSS bidirectionals, genes, and tREs across increasing replicate numbers (colors) and quantiles (Q1-Q10) according to average counts (log2). Due to the different mean distributions across the feature types, the quantiles do not represent the same count levels. The grey box represents a section of features with comparable average counts (around 5.5-7). Shade of green reflects replicate numbers per condition.

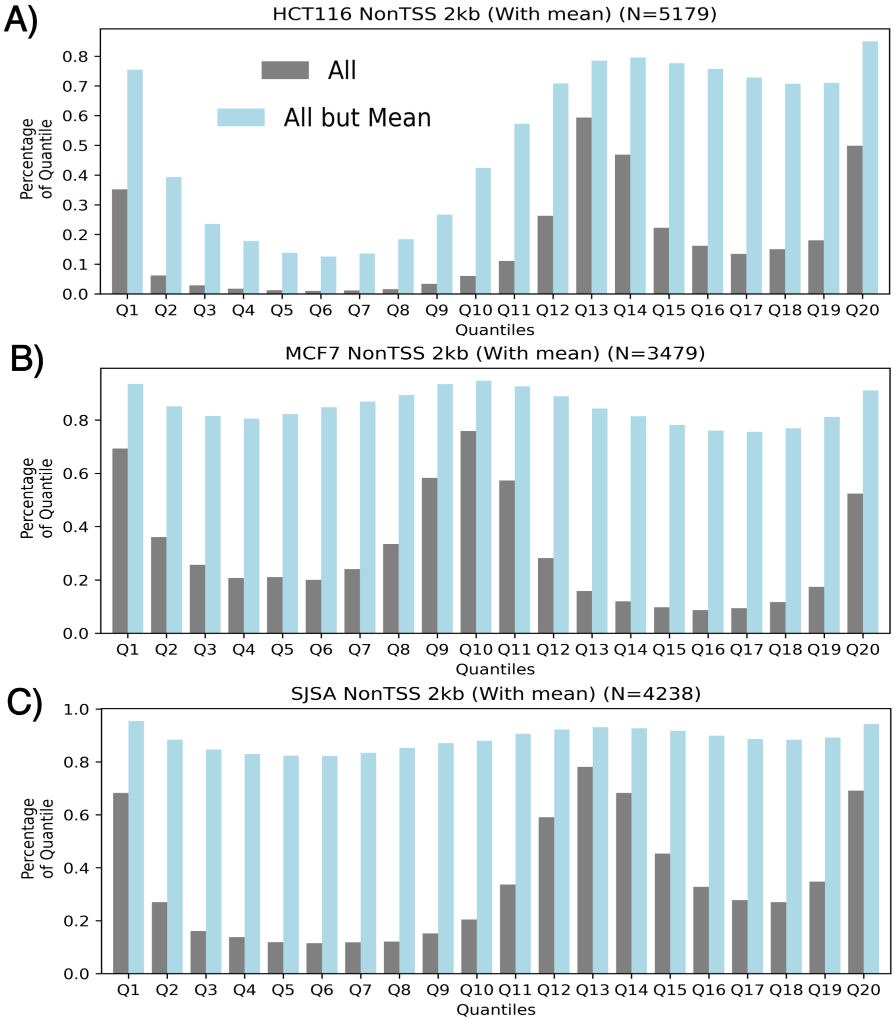


Figure S5: **The features with highest statistical confidence for differential transcription are consistent across classic differential expression tools for all celltypes.** The percentage of tREs within the same ranked quantile according to all tested tool-parameter combinations (grey) or all excluding combinations using the Mean-based dispersion estimation (light blue) for (A) HCT116, (B) MCF7, and (C) SJSA. tREs are ranked according to direction of change and adjusted p-value where the poles have the greatest statistical significance and the middle tREs have little to no change.

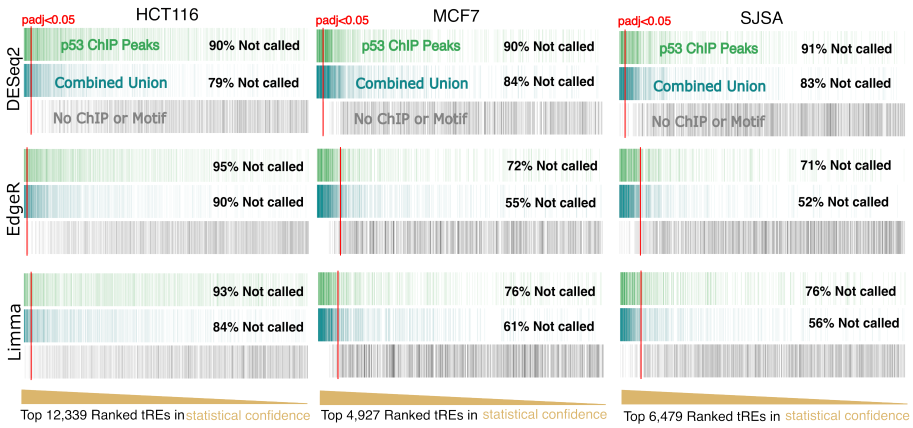


Figure S6: **True positives are often mixed with statistical confidence of expected true negatives, regardless of cell type and tool-based rankings.** The top tREs with positive log fold change for p53 ranked according to adjusted p-values values (proxy for statistical confidence) for each cell type. The tREs overlapping p53 ChIP peaks for the appropriate cell type are colored green (top), called when using all cell types as replicates ("Combined Union") are colored turquoise (middle), and those with no clear linkage to Nutlin-3a or P53 are colored grey (bottom). A red line corresponds to the position at which all tREs to the left are called significant at p-adjusted value cutoff of 0.05 (padj$<$ 0.05). HCT116 is expected to have very conservative classic results for EdgeR and Limma due to one of the samples having significantly lower overall transcription levels than the rest of the samples [2].

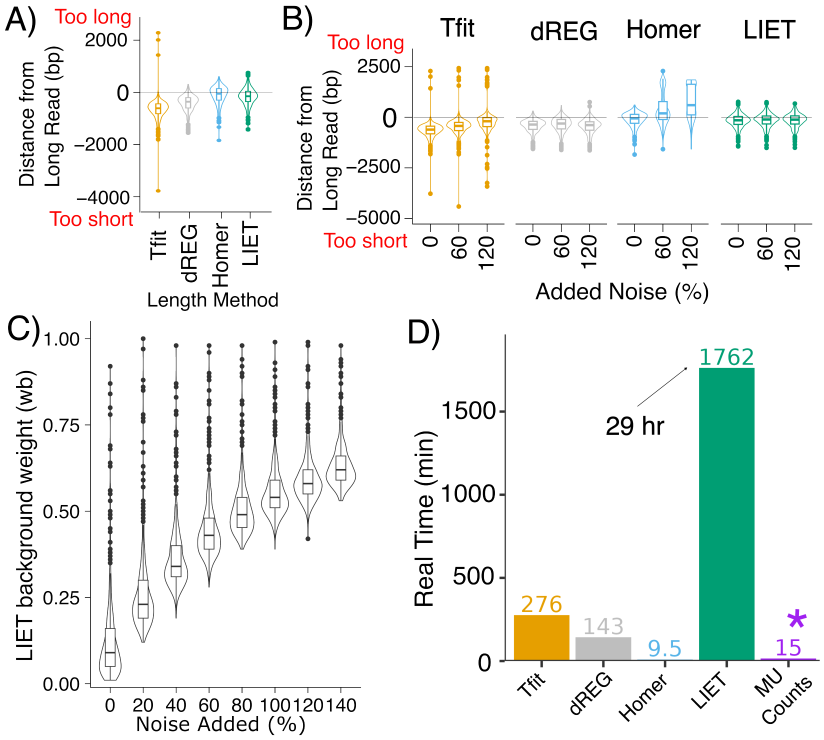


Figure S7: **Despite taking the longest, tRE-adapted LIET consistently produces lower error in tRE length after considering overlapping transcription.** Results of the four tRE identification methods (Tfit (yellow), dREG (grey), Homer (blue), LIET (green) on length prediction across 411 tREs, both without (**A**) and with (**B**) noise added. Distance from Long Read serves as a proxy for length prediction error (details in Supplementary Methods). A negative value refers to the prediction being too short, and a positive to the prediction being too long. Results for A and B that consider all twelve methods (as described in Supplementary Methods) and all noise levels can be found at (**github:**/Length_Bench/Comparison/Compare_Lengths.ipynb). All results correspond to the short-read data from SRA SRR4454567. Similar results were found when calculating consensus lengths from tools. **C.** LIET background weight (Bayesian prior (posterior estimated graphed here)) effectively captures noise. Results from other samples and LIET-model adaptations can be found in the same notebook above. **D.** Real time taken to run each of the length-predicting methods on one sample, or in the case of mu-Counts in all samples at once. Homer, the fastest approach for a single sample, takes 25 minutes to run on all samples at once.

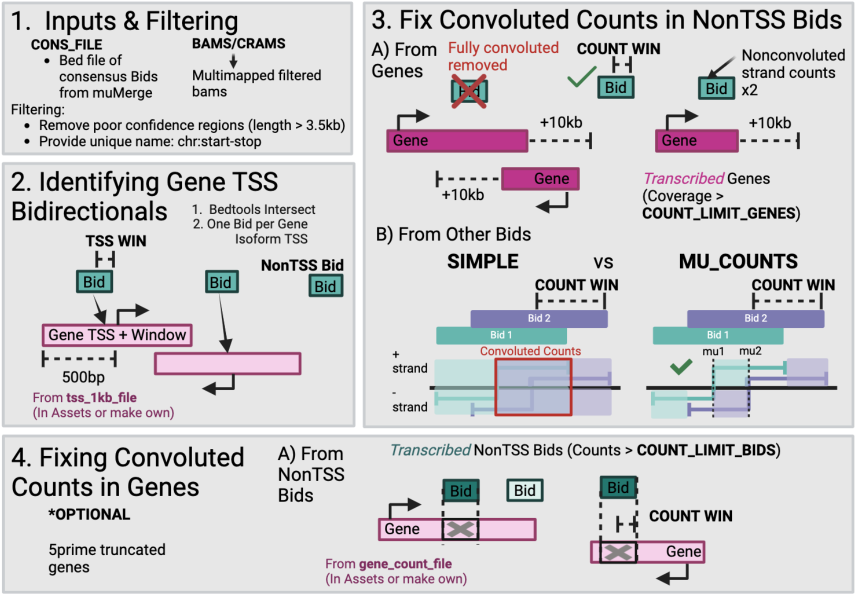


Figure S8: **Visual representation of the Mu_Counts pipeline.** Full description in Supplemental Methods **Step 1**-Filtering Inputs. We remove bidirectionals that are likely called due to technical noise and filter bams/crams to remove multi-mapped reads. This produces consensus files with the widths needed for future analyses and unique names. **Step 2**-Identifying Gene TSS Bidirectionals. Bidirectionals coordinating to the transcription start site (TSS) of genes are identified by overlapping bidirectionals with a small window (parameter TSS_WIN - default 25bp) with 1kb regions around gene TSSs (parameter tss_1kb_file, also available for GR38.p14). **Step 3A**-Addressing overlapping transcription from genes. Gene bodies considered transcribed (COUNT_LIMIT_GENES=70% isoform is covered with reads) along with the region 10kb downstream of said isoform are overlapped with the nonTSS bidirectionals (tREs). Any nonTSS bidirectional overlapping active gene transcription on both strands is removed since deconvolution cannot confidently occur. NonTSS bidirectionals overlapping gene transcription on one strand have counts replaced with those from the on-convoluted strand, doubled. **Step 3B**-Addressing overlapping transcription from bidirectionals with other bidirectionals (Mu_Counts). If the parameter COUNT_WIN for a tRE means the tRE is now overlapping with another, the neighborhing tREs will be counted so that the maximum distance of each RNA is the $\mu$ of the nearest neighboring tRE. RNAs for tREs are then counted separately according to the strand (e.g. counts on positive/negative strand for Bid 2's positive/negative RNA are combined). **Step 4**-Fixing gene counts due to overlapping bidirectionals. The pipeline optionally removes the regions of bidirectionals with counts above parameter COUNT_LIMIT_BIDS.

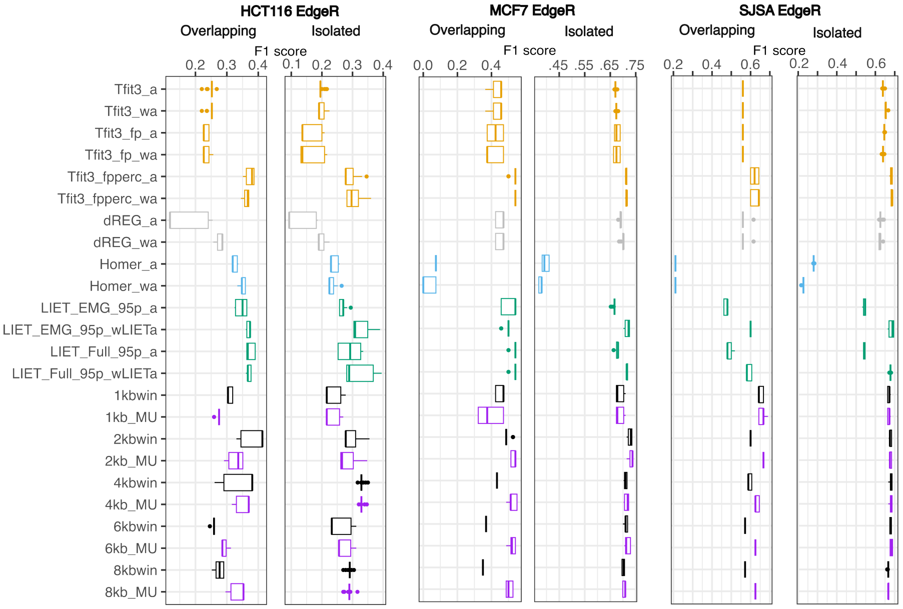


Figure S9: **tRE-focused LIET and Mu_Counts both show the highest F1 scores for overlapping and isolated tREs across all cell types.** F1 scores using p53 ChIP-peaks as the truth set for HCT116, MCF7, and SJSA when using EdgeR and counts from windows defined by multiple different methods. Methods colored as Figure S7. Fixed_win refers to a fixed window and Fixed_win_mu (and suffix win) refers to the Mu_Counts (and suffix MU) pipeline being used with the provided initial fixed window. Average is noted by _a, weighted average (based on counts) is noted by _wa. Tfit includes fp and fpperc where the footprint is added to Tfit length, with or without the 95th percentile of the EMG, respectively. EMG_95p means that the 95th percentile of LIET using just the EMG was used. wLIET means that the weighted average was used with weights based on $wLIET = 1 - w_b$ ($w_b$ =background weight in LIET). Details on other methods can be found in Supplementary Methods section. All comparisons (e.g. cell types and platforms) can be found at (**github:**/Bench_DE/Length_DE/p53_Len_Compare_Vis.ipynb).

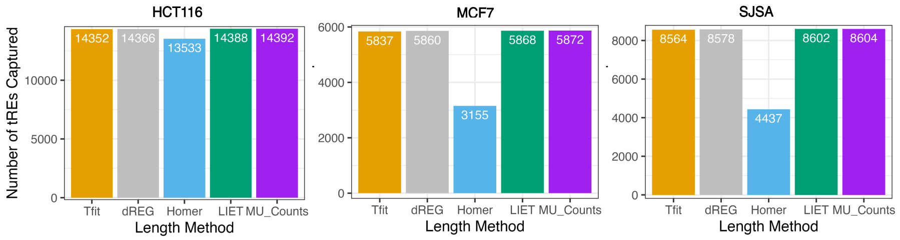


Figure S10: **Homer provides length for significantly less tREs compared to its counterparts.** The number of tREs used for length-based differential transcription assessment whose lengths could be estimated by the relevant methods: Tfit (Tfit with 3′ bedgraphs), dREG, Homer, LIET (with 1- $w_b$ used for consensus calculation in LIET (details in Supplemental Methods)), and Mu_Counts (so all). The total tREs are based on the consensus tREs determined as described in Supplementary Methods, so that all methods are considering the same "universe" of tREs. Importantly, LIET uses pre-annotated search positions on which to model tREs while Homer cannot take predefined regions on which to consider its model. Therefore, LIET and Mu_Counts have a huge advantage to consider any pre-specified tREs (as defined by tRE identification tools like Tfit and dREG).

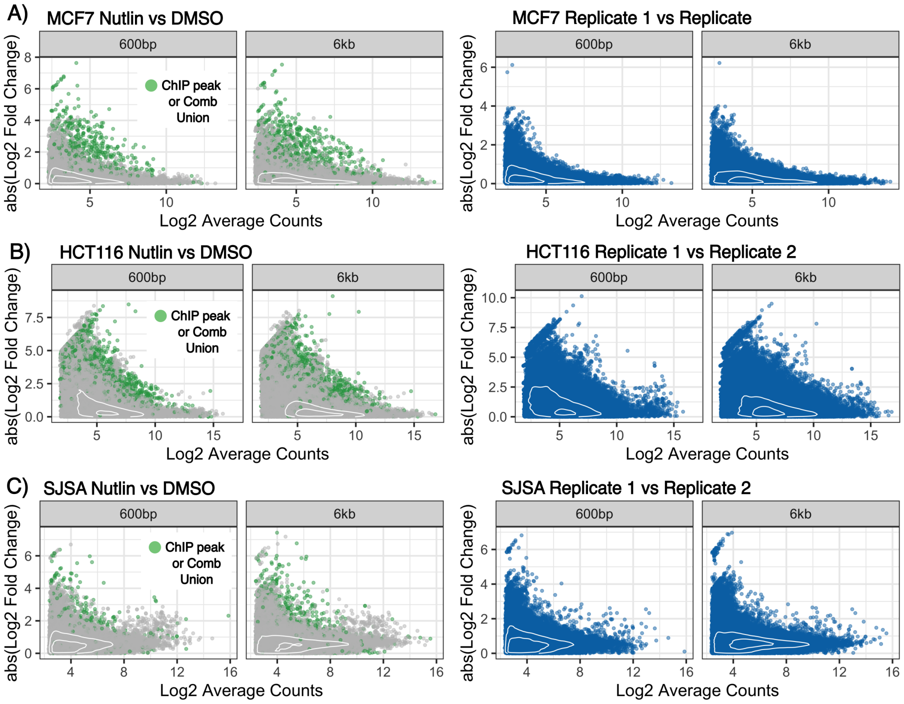


Figure S11: **Mean-dispersion trends remain similar, leading to low statistical confidence for p53 data, despite length correction with Mu_Counts.** Absolute log fold change of tREs between Nutlin-3a and DMSO (left, green/grey) and biological replicates (right, blue) with tREs that are supported by ChIP peaks or with combined cell types ("Comb Union") are highlighted in green. Results are considered for A) MCF7, B) HCT116, and C) SJSA when using counts from 600bp fixed windows (left) and 6kb Mu_Counts (right).

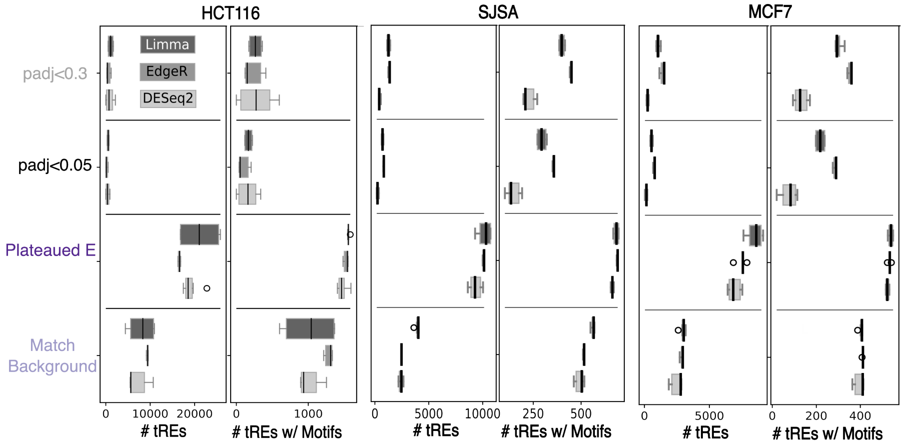


Figure S12: **Leading Edge positions are consistent across ranking platforms.** Boxplots of leading edge positions for both leading edge methods and classic statistical tools (padj < 0.3 and padj < 0.05) when considering all tREs or just those with the corresponding TF motifs within 1.5kb of the tRE $\mu$s (midpoints). The variability of Limma-Voom vs Limma-Trend leading edge results in HCT116 is not observed with any other cell type or condition.

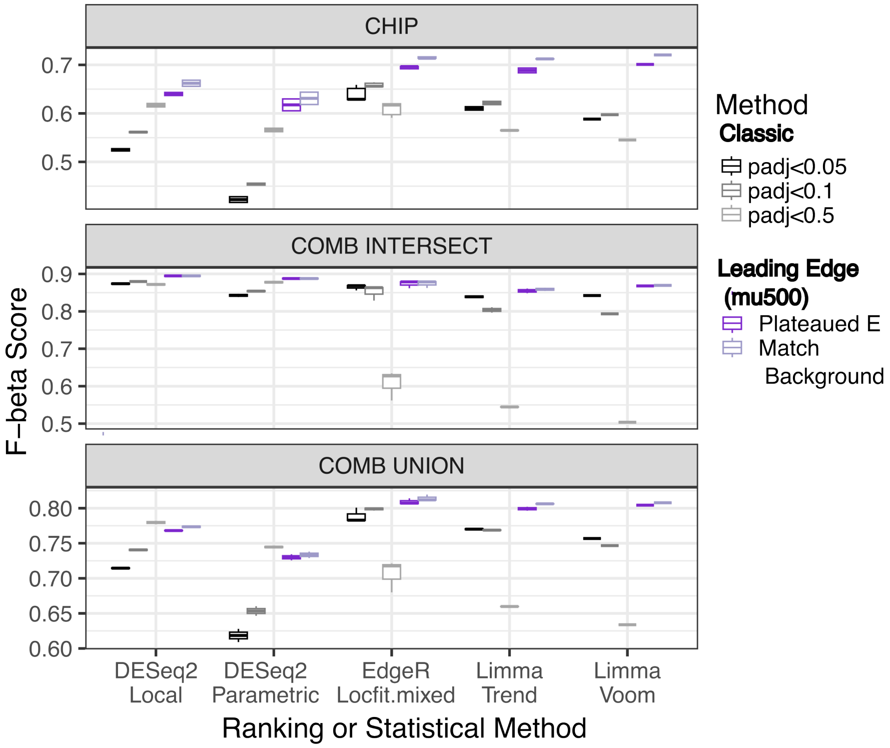


Figure S13: **The leading edge methods confer improved balance of precision and recall based on F1-beta scores.** F1-beta scores of p53 responsive tREs based on their proximity to corresponding p53 ChIP peaks. These results are from SJSA using Mu_Counts with a max window size of 2kb since they were well-representative of all results. Other cell type and window size results are comparable but can be found at (**github:**/Compare_LE/p53_Compare_LE_stats.ipynb). Graphs with recall and precision mapped as scatter plots are also available at the same notebook. Due to the extreme conservativeness of DESeq2, adjusted p-values of 0.5 were occasionally able to allow a small increase in recall compared to leading edges while maintaining precision above 0.9.

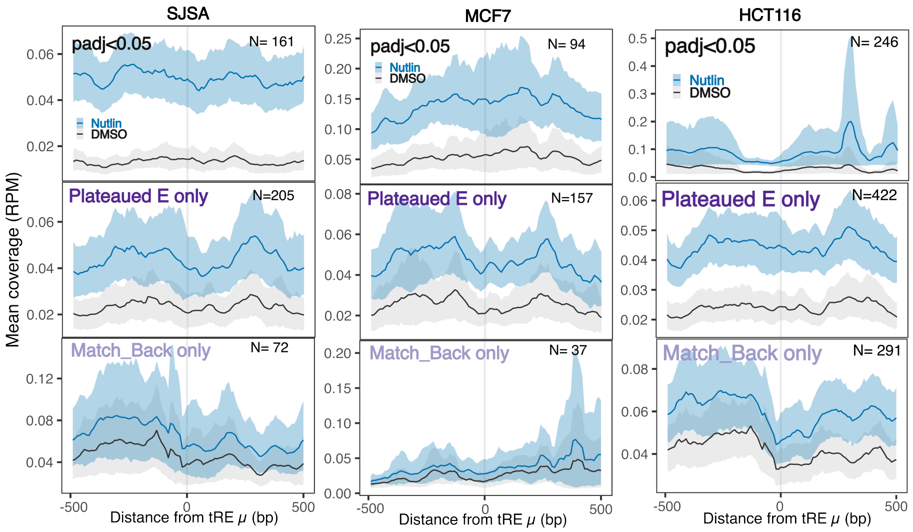


Figure S14: **Metaplots of p53 reveal Plateaued Enrichment leading edge confidently captures tREs with changes in transcription.** Metaplots of the tREs normalized mean coverage (Reads Per Million) found with classic statistical approaches, or leading edge methods with motifs within 500bp of the tRE centers. The lines (DMSO: grey, Nutlin-3a: blue) refer to the mean whereas highlighted regions are the 95% confidence interval of read distribution for tREs. Results shown here are based on 2kbmin Mu_Counts and DESeq2: Ratio_LRT_Local but are representative of those from other classic statistical approaches.

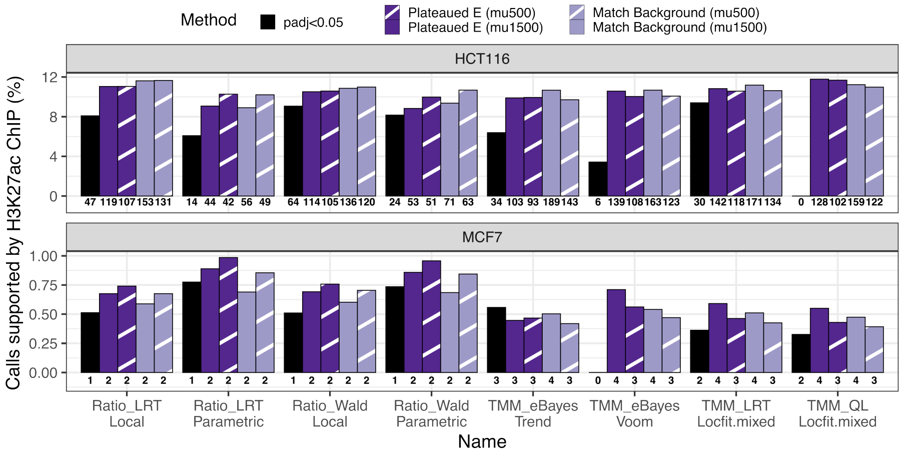


Figure S15: **Leading edge calls have higher enrichment of H3K27ac support.** Percentage of calls for HCT116 (top) and MCF7 (bottom)supported by ChIP peaks. (Note H3K27ac unavailable for Nutlin-3a in SJSA cells). A H3K27ac peak must be only called after Nutlin-3a has been added to media (1hr for HCT116 and 2.5hr for MCF7). p05 refers to the classic statistical approach cutoff for various methods (column labels), Plateaued E for adding tREs within the "Plateaued Enrichment" leading edge that have a p53 motif within 500bp (mu500) or 1.5kb (mu1500) of their midpoint. Same for "Match Background" leading edge.

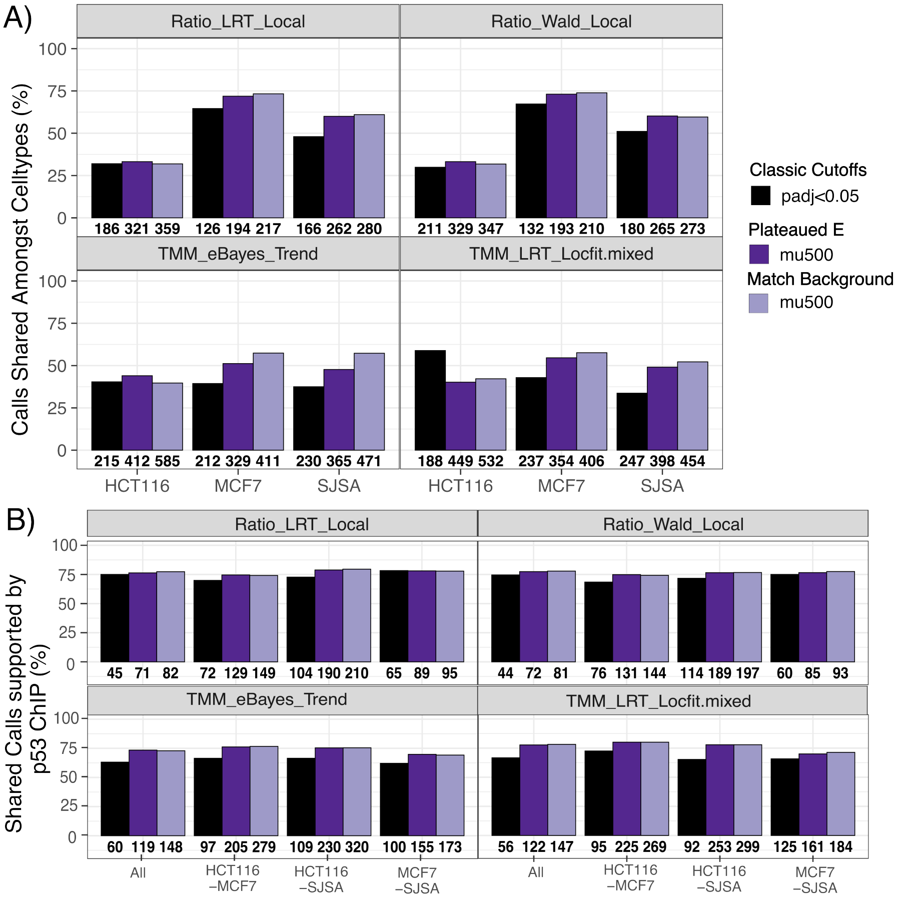


Figure S16: **Despite leading edge calling more p53-responsive tREs shared across cell types, these calls are generally supported by p53 ChIP peaks A.** Percentage of tREs for each cell type that are called in at least one other celltype (i.e. shared) according to classic tools (padj<0.05) (noted in titles - e.g. Ratio_LRT_Local), or leading edge methods including calls with p53 motifs within 500bp of tRE midpoints (Plateaued E (mu500) and Match Background (mu500)) **B.** Percentage of calls shared between all celltypes or pairwise combinations that overlap p53 ChIP peaks shared across cell types. In all cases, the bottom numbers refer to the N of the bars while the y values graphed are the percentages of the total calls within each method. Results from all classic tool-parameter combinations as well as leading edge with tREs containing motifs within 1.5kb can be found at (**github:**/Bench_DE/Compare_LE/p53_Compare_Celltypes.ipynb)

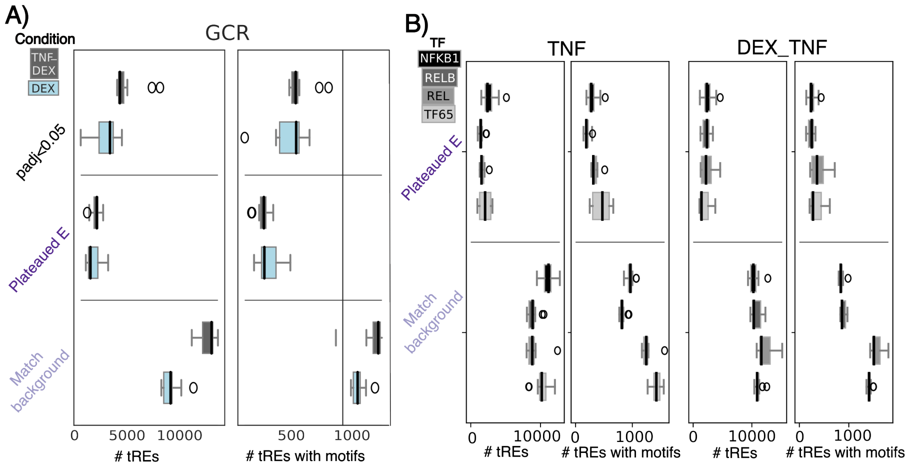


Figure S17: **Leading Edge positions for GR and NF$\kappa$B TFs are consistent across ranking methods.** Boxplots of leading edge positions in A) dexamethasone (DEX), B) TNF, or C) dexamethasone and TNF (DEX_TNF) when considering all tREs (left) or just those with the corresponding TF motifs (GR, NFKB1, RELB, REL, TF65) within 1.5kb of the tRE $\mu$s (right). Boxplots represent leading edge or classic statistical methods with padj<0.05.

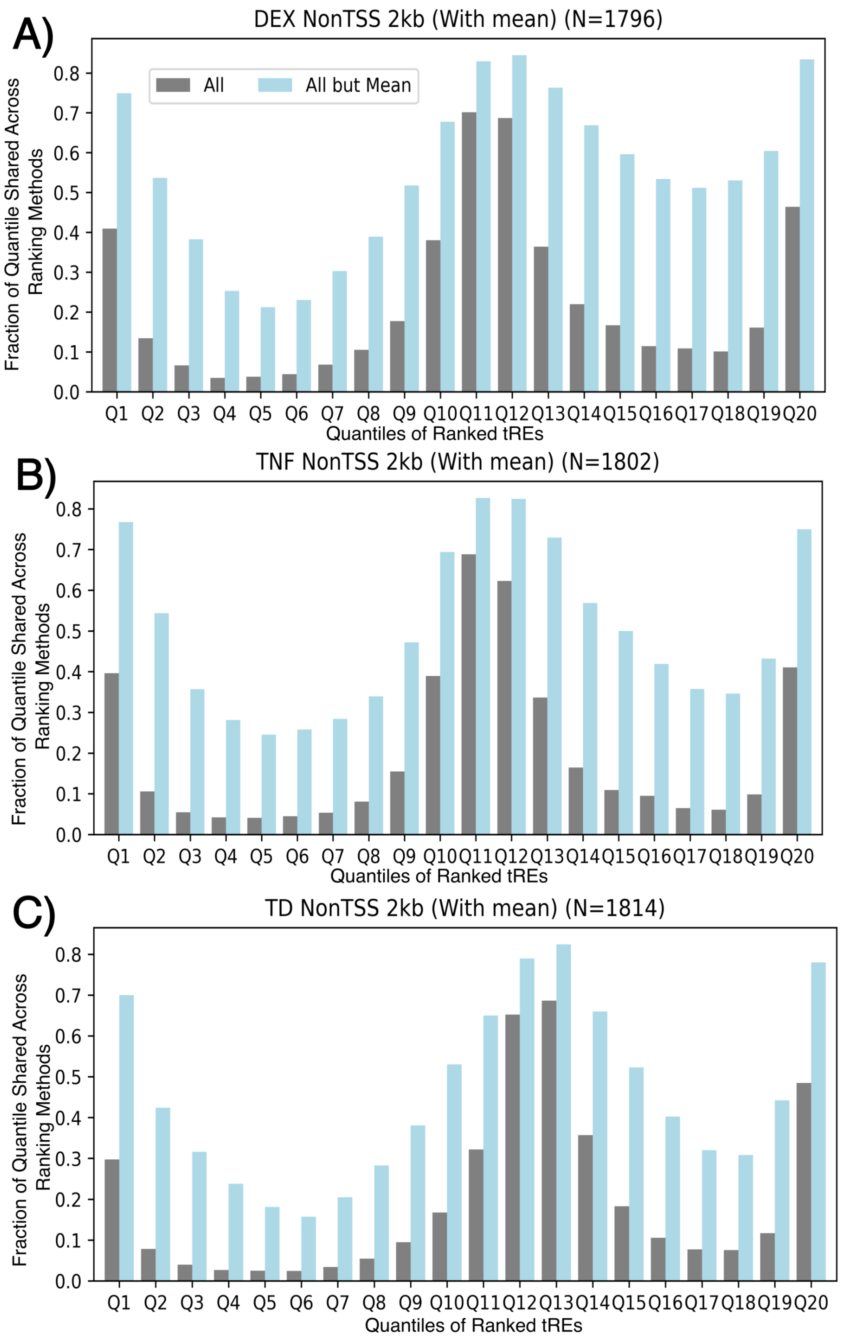


Figure S18: **tREs ranked with the greatest statistical confidence of change from classic tools for are consistent for TNF/DEX perturbations.** The percentage of tREs within the same ranked quantile of (A) DEX, (B) TNF, and (C) both TNF and DEX according to all tested tool-parameter combinations (grey) or all excluding combinations using the Mean-based dispersion estimation (light blue). tREs are ranked according to direction of change and adjusted p-value (poles have greatest statistical significance and middle has little to no change).

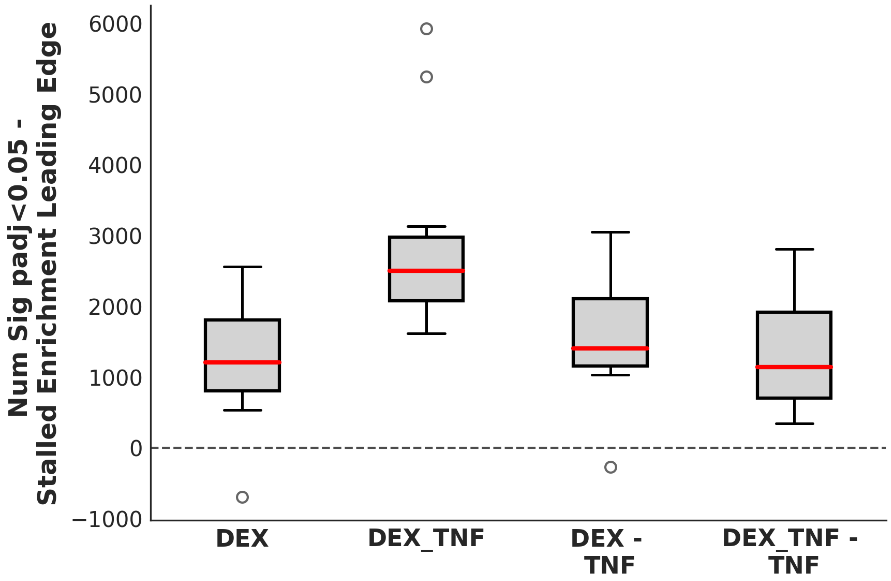


Figure S19: **Unlike with Nutlin-3a and p53, classic tools call more tREs significant than the GR leading edge for DEX and TNF perturbed cells.** Boxplots of the difference between the number of significant calls according to adjusted p-values $< 0.05$ and all tREs (regardless of motif) within the Plateaued Enrichment leading edge. Dexamethasone (DEX), dexamethasone with TNF (DEX_TNF), and these conditions with the TNF only significant (adjp $< 0.05$) calls removed (-TNF). The only case where the classic statistical approach does not call more significant tREs is for Limma (eBayes significance test and Trend dispersion estimation).

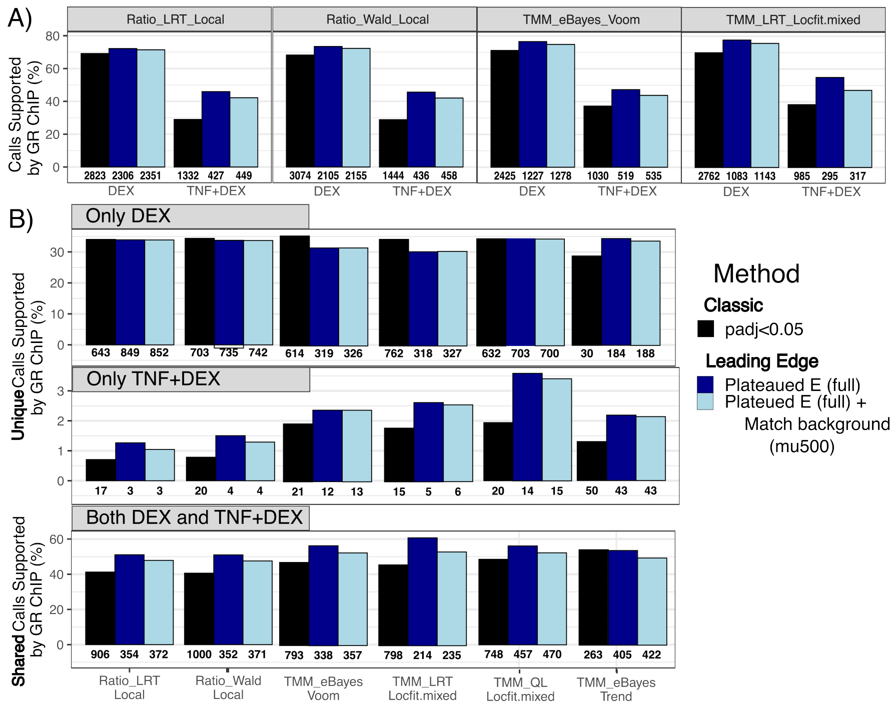


Figure S20: **Leading Edge based calls for GR for are equally or more enriched in GR ChIP peaks than calls with classic tools. A.** The percentage of calls supported by ChIP-seq peaks of GR for cells treated with dexamethasone (DEX) or both dexamethasone and TNF (TNF+DEX). padj<0.05 refers to the classic approach. Plateaued E (full) means all calls within the Plateaued E leading-edge are considered. Plateaued E (full) + Match Background (mu500) adds tREs within the Match Background leading-edge that have a GR motif within 500bp of their midpoints. Tool-parameter combinations are representative of all results. **B.** The percentage of calls considered either unique to DEX or TNF+DEX perturbed cells and percentage of calls considered shared by both perturbations supported by equivalent comparisons with GR ChIP. Tool-parameter combinations are representative of all results. Calls from leading edge with the classic approach and motif calls (e.g. mu500) are not shown since they give almost equivalent results to the classic statistical approach. Full results including these and all tool-parameter combinations can be found at (**github:**/Bench_DE/Compare_LE/GR_Compare_Conditions.ipynb).

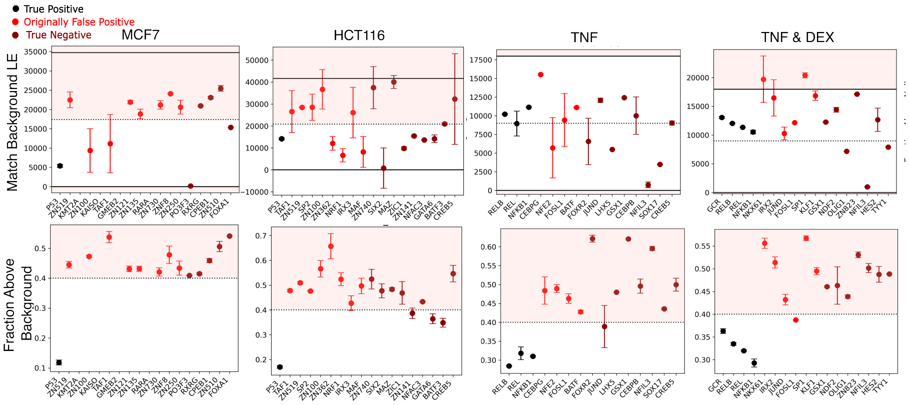


Figure S21: **Leading-edge related values serve as robust secondary metrics of false positive TFEA calls.** TFs are color-coded as known expected calls (black), improperly called significant (light red), or properly called non-significant enriched (dark red). Dots indicate the number of tREs within a Match Background leading edge (top) or fraction of tREs with cumulative enrichment scores above that expected from background (bottom). The quarter point of tREs and below 0 (top) or fraction above 0.4 (bottom) are highlighted. HCT116 and MCF7 refer to Nutlin-3a (p53) datasets for these celltypes. TNF and TNF & DEX refer to lung cells perturbed with TNF or both TNF and dexamethasone. Only these are shown for brevity; other celltypes and perturbations can be found at (**github:**/Improving_FP_calls/LE_FP.ipynb).

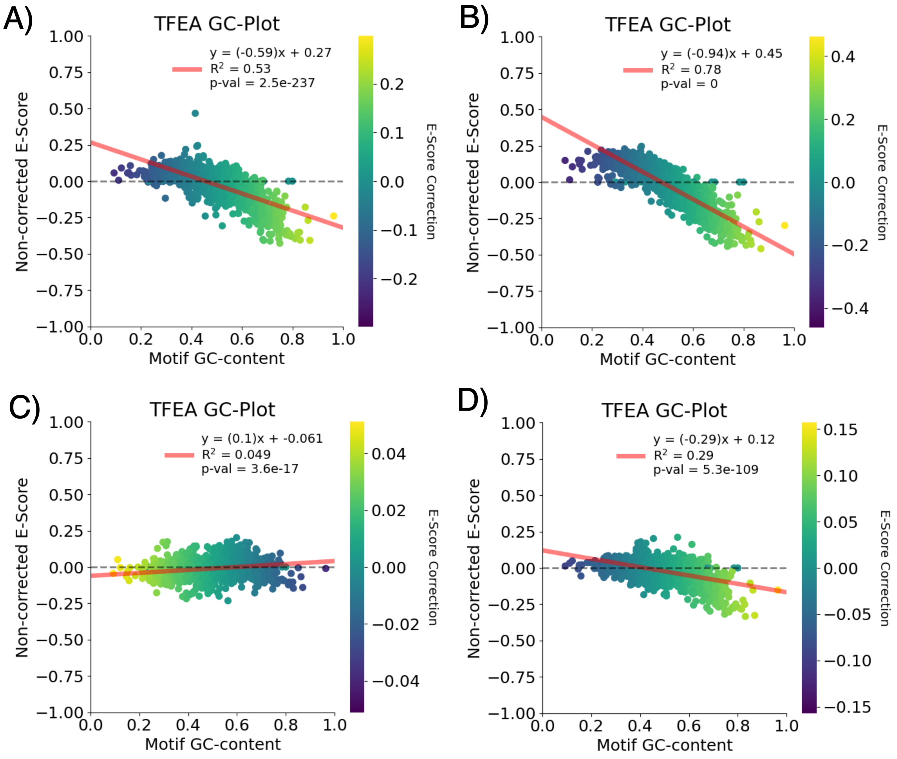


Figure S22: **PRO-seq and to a lesser extent ATAC-seq data for woodsmoke particles has strong GC-bias.** GC-correction graphs from TFEA for A) PRO-seq 30min, B) PRO-seq 120min, C) ATAC-seq 30min, D) ATAC-seq 120min. Each dot represents a TF with its original score (non-corrected E-score, y-axis) against its GC-content (y-axis) and colored by the new E-score after correction.

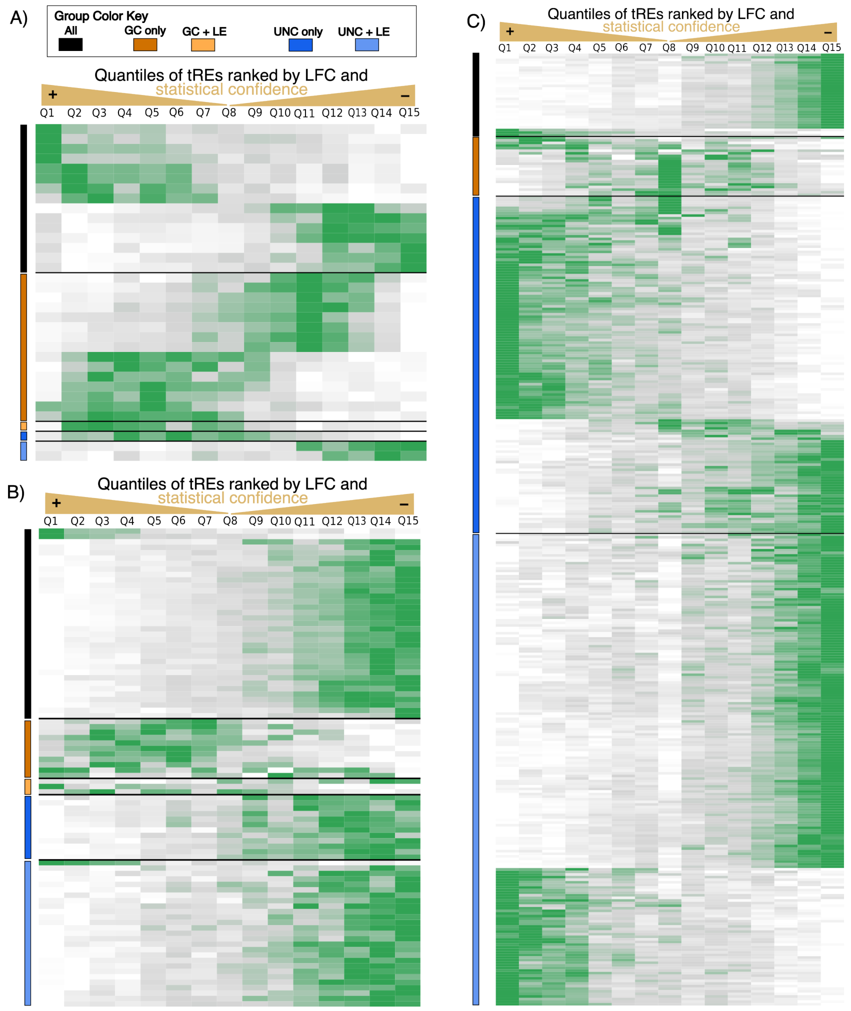


Figure S23: **Leading edge metrics successfully remove false positive TF calls in both ATAC and PRO-seq, regardless of GC correction** Quantile enrichment plots for A) ATAC-seq WSP at 30min vs Veh, B) ATAC-seq WSP 120min vs Veh, and C) PRO-seq WSP 120min vs Veh. Transcription factor calls are split into All (black, LE metrics support, GC-corrected and uncorrected adjusted p-values < 0.001), GC only (red, GC-corrected but not uncorrected adjusted p-value < 0.001 and not supported by LE-metrics), GC-LE (orange, supported by GC-correction and LE-metrics but not uncorrected adjusted p-value), UNC only (dark blue, supported by only uncorrected adjusted p-value), UNC+LE (light blue, supported by only uncorrected adjusted p-value and LE-metrics). Slope at quantiles of tREs (each quantile has 2,944 tREs). Ns: ATAC-seq 30min All=15, GC only=15, GC+LE=1, UNC only=1, UNC+LE=2; ATAC-seq 120min All=39, GC only=11, GC+LE=3, UNC only=12, UNC+LE=29; PRO-seq 120min All=33, GC only=24, GC+LE=0, UNC only=137, UNC+LE=207.

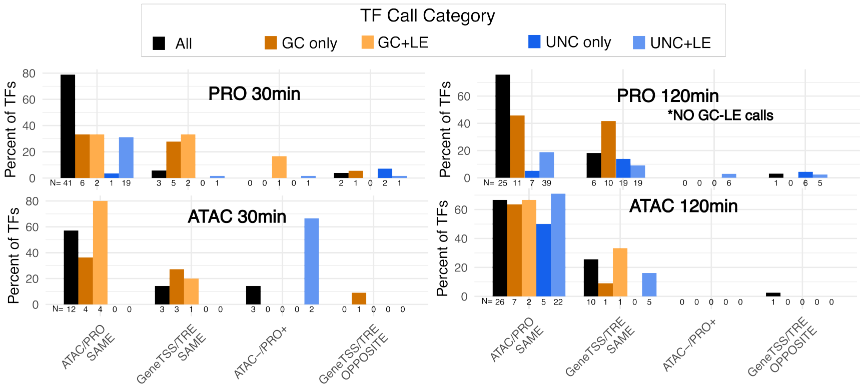


Figure S24: **Direction of TF significant calls between ATAC and PRO or gene TSS bidirectionals and tREs was mostly enriched when including leading-edge focused metrics.** Percentage of TFs from each of the TF call categories that have enrichment scores going the same direction in ATAC-seq and PRO-seq (ATAC/PRO SAME) or with gene TSS bidirectionals and tREs (GeneTSS/TRE SAME), or suggest closing of chromatin but increased transcription (ATAC-/PRO+) or opposite directions between gene TSS bidirectionals and tREs (GeneTSS/TRE OPPOSITE). All refers to TFs called significant by GC-correction, no correction, and LE metrics. GC only and GC+LE refer to TFs called with GC-corrected but not uncorrected significance, and called without or with LE metric support, respectively. UNC only and UNC+LE refer to TFs called with uncorrected but not GC-corrected significance, and called without or with LE metric support, respectively. PRO-seq 120 minutes did not have any TFs supported by both GC-correction and Leading edge metrics (NO GC-LE calls).

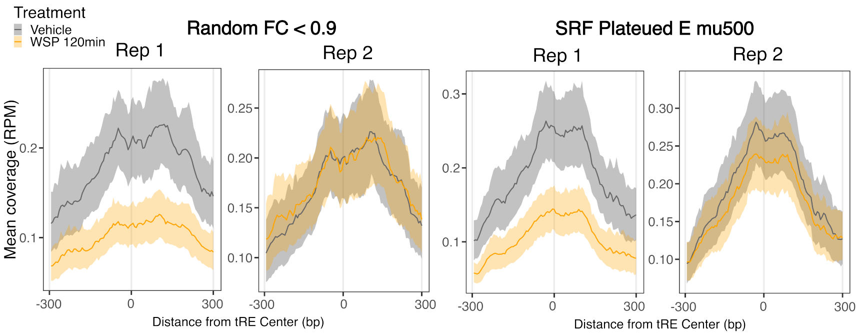


Figure S25: **Leading edge tREs show decreased accessibility in both replicates while the same number of random tREs with fold changes below 0.9 do not.** Mean reads per-million of the two ATAC-seq vehicle and 120 minute WSP replicates of tREs randomly selected from tREs with DESeq2 fold changes below 0.9 (Random FC $<$ 0.9) vs those with an SRF motif within the Plateaued E Leading edge (SRF Plateaued E mu500) (both N=98). Only 1 tRE is found in both sections. Results for SRF Match Background mu500 (with its own random set) look almost identical. The tREs that have FC below 0.9 but are outside the Plateaued E LE show lower decreased accessibility with overlaps of confidence intervals for combined replicates, unlike the tREs of the same number just barely within the leading edge. These graphs can be found at (**github:**/WSP/Graph_WSP_metaplots.ipynb).

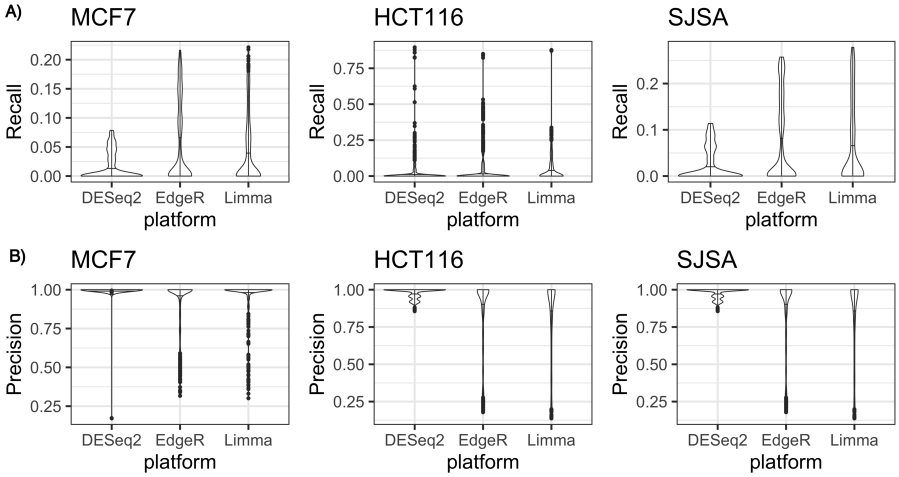


Figure S26: **AUC-PR metrics would be inappropriate due to variable ranges of precision and recall across tools and parameter combinations for p53 based analysis.** Precision and recall for the three tools (DESeq2, EdgeR, and Limma) were tested using a broad range of parameters and p-value cutoffs (see Supplementary Methods), and results are shown as boxplots. MCF7, HCT116, and SJSA refer to the cell types considered. The true positive set used was based on ChIP identified p53 calls (details in Supplementary Methods).

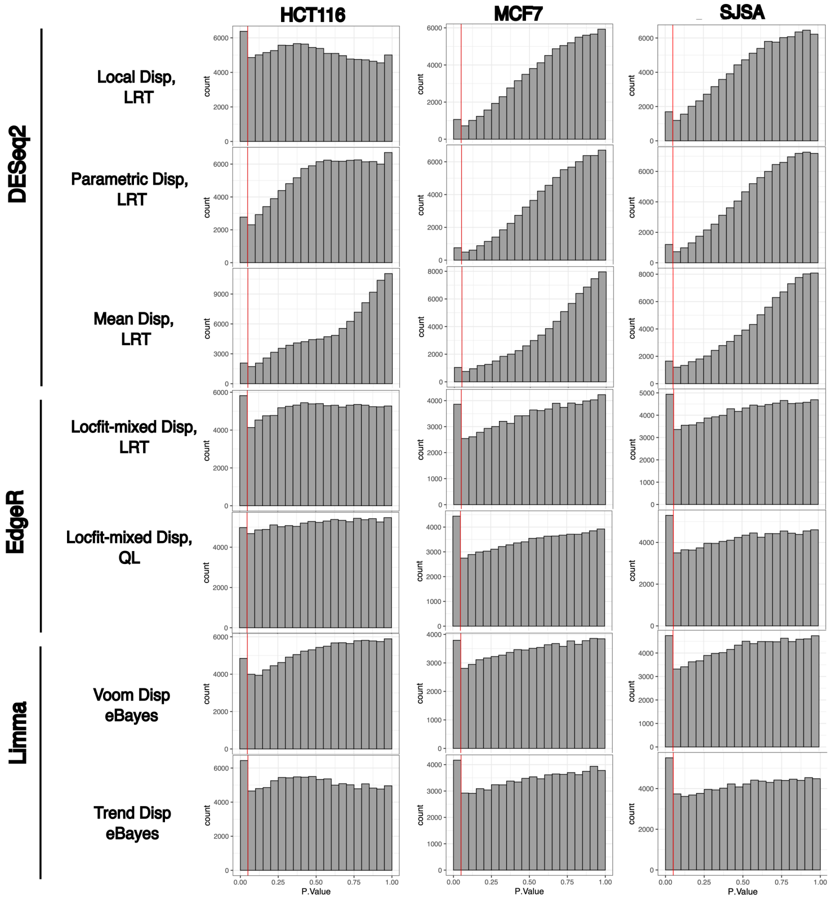


Figure S27: **Distribution of p-value scores (non-adjusted) reflects unclean analysis particularly for DESeq2, and mean-dispersion estimation.** The red line indicates the 0.05 position (common p-value cutoff). Built-in normalization factors (Ratio for DESeq2, TMM for EdgeR and Limma) were used here but use of virtual-spike in normalization factors show almost identical results (data not shown) [12]. DESeq2 Mean Dispersion (Disp) shows bumps in p-values common with poor dispersion calling. EdgeR and Limma showed the strongest consistency in a peak at the 0-0.05 p-value position with a generally uniform distribution after. All outputs were from using Mu_Counts with a 2kb fixed window to ensure biased counting approaches did not explain comparisons. To simplify visualization, only one of tested combinations with almost identical results are shown (e.g. DESeq2 Wald vs LRT, QL-Robust vs QL).

#### References

[1] Federico Abascal, Reyes Acosta, Nicholas J. Addleman, Jessika Adrian, Veena Afzal, Rizi Ai, Bronwen Aken, Jennifer A. Akiyama, Omar Al Jammal, Henry Amrhein, Stacie M. Anderson, Gregory R. Andrews, Igor Antoshechkin, Kristin G. Ardlie, Joel Armstrong, Matthew Astley, Budhaditya Banerjee, Amira A. Barkal, If H. A. Barnes, Iros Barozzi, Daniel Barrell, Gemma Barson, Daniel Bates, Ulugbek K. Baymuradov, Cassandra Bazile, Michael A. Beer, Samantha Beik, M. A. Bender, Ruth Bennett, Louis Philip Benoit Bouvrette, Bradley E. Bernstein, Andrew Berry, Anand Bhaskar, Alexandra Bignell, Steven M. Blue, David M. Bodine, Carles Boix, Nathan Boley, Tyler Borrman, Beatrice Borsari, Alan P. Boyle, Laurel A. Brandsmeier, Alessandra Breschi, Emery H. Bresnick, Jason A. Brooks, Michael Buckley, Christopher B. Burge, Rachel Byron, Eileen Cahill, Lingling Cai, Lulu Cao, Mark Carty, Rosa G. Castanon, Andres Castillo, Hassan Chaib, Esther T. Chan, Daniel R. Chee, Sora Chee, Hao Chen, Huaming Chen, Jia-Yu Chen, Songjie Chen, J. Michael Cherry, Surya B. Chhetri, Jyoti S. Choudhary, Jacqueline Chrast, Dongjun Chung, Declan Clarke, Neal A. L. Cody, Candice J. Coppola, Julie Coursen, Anthony M. D'Ippolito, Stephen Dalton, Cassidy Danyko, Claire Davidson, Jose Davila-Velderrain, Carrie A. Davis, Job Dekker, Alden Deran, Gilberto DeSalvo, Gloria Despacio-Reyes, Colin N. Dewey, Diane E. Dickel, Morgan Diegel, Mark Diekhans, Vishnu Dileep, Bo Ding, Sarah Djebali, Alexander Dobin, Daniel Dominguez, Sarah Donaldson, Jorg Drenkow, Timothy R. Dreszer, Yotam Drier, Michael O. Duff, Douglass Dunn, Catharine Eastman, Joseph R. Ecker, Matthew D. Edwards, Nicole El-Ali, Shaimae I. Elhajjajy, Keri Elkins, Andrew Emili, Charles B. Epstein, Rachel C. Evans, Iakes Ezkurdia, Kaili Fan, Peggy J. Farnham, Nina P. Farrell, Elise A. Feingold, Anne-Maud Ferreira, Katherine Fisher-Aylor, Stephen Fitzgerald, Paul Flicek, Chuan Sheng Foo, Kevin Fortier, Adam Frankish, Peter Freese, Shaliu Fu, Xiang-Dong Fu, Yu Fu, Yoko Fukuda-Yuzawa, Mariateresa Fulciniti, Alister P. W. Funnell, Idan Gabdank, Timur Galeev, Mingshi Gao, Carlos Garcia Giron, Tyler H. Garvin, Chelsea Anne Gelboin-Burkhart, Grigorios Georgolopoulos, Mark B. Gerstein, Belinda M. Giardine, David K. Gifford, David M. Gilbert, Daniel A. Gilchrist, Shawn Gillespie, Thomas R. Gingeras, Peng Gong, Alvaro Gonzalez, Jose M. Gonzalez, Peter Good, Alon Goren, David U.

Gorkin, Brenton R. Graveley, Michael Gray, Jack F. Greenblatt, Ed Griffiths, Mark T. Groudine, Fabian Grubert, Mengting Gu, Roderic Guigó, Hongbo Guo, Yu Guo, Yuchun Guo, Gamze Gursoy, Maria Gutierrez-Arcelus, Jessica Halow, Ross C. Hardison, Matthew Hardy, Manoj Hariharan, Arif Harmanci, Anne Harrington, Jennifer L. Harrow, Tatsunori B. Hashimoto, Richard D. Hasz, Meital Hatan, Eric Haugen, James E. Hayes, Peng He, Yupeng He, Nastaran Heidari, David Hendrickson, Elisabeth F. Heuston, Jason A. Hilton, Benjamin C. Hitz, Abigail Hochman, Cory Holgren, Lei Hou, Shuyu Hou, Yun-Hua E. Hsiao, Shanna Hsu, Hui Huang, Tim J. Hubbard, Jack Huey, Timothy R. Hughes, Toby Hunt, Sean Ibarrientos, Robbyn Issner, Mineo Iwata, Osagie Izuogu, Tommi Jaakkola, Nader Jameel, Camden Jansen, Lixia Jiang, Peng Jiang, Audra Johnson, Rory Johnson, Irwin Jungreis, Madhura Kadaba, Maya Kasowski, Mary Kasparian, Momoe Kato, Rajinder Kaul, Trupti Kawli, Michael Kay, Judith C. Keen, Sunduz Keles, Cheryl A. Keller, David Kelley, Manolis Kellis, Pouya Kheradpour, Daniel Sunwook Kim, Anthony Kirilusha, Robert J. Klein, Birgit Knoechel, Samantha Kuan, Michael J. Kulik, Sushant Kumar, Anshul Kundaje, Tanya Kutyavin, Julien Lagarde, Bryan R. Lajoie, Nicole J. Lambert, John Lazar, Ah Young Lee, Donghoon Lee, Elizabeth Lee, Jin Wook Lee, Kristen Lee, Christina S. Leslie, Shawn Levy, Bin Li, Hairi Li, Nan Li, Shantao Li, Xiangrui Li, Yang I. Li, Ying Li, Yining Li, Yue Li, Jin Lian, Maxwell W. Libbrecht, Shin Lin, Yiing Lin, Dianbo Liu, Jason Liu, Peng Liu, Tingting Liu, X. Shirley Liu, Yan Liu, Yaping Liu, Maria Long, Shaoke Lou, Jane Loveland, Aiping Lu, Yuheng Lu, Eric Lécuyer, Lijia Ma, Mark Mackiewicz, Brandon J. Mannion, Michael Mannstadt, Deepa Manthravadi, Georgi K. Marinov, Fergal J. Martin, Eugenio Mattei, Kenneth McCue, Megan McEown, Graham McVicker, Sarah K. Meadows, Alex Meissner, Eric M. Mendenhall, Christopher L. Messer, Wouter Meuleman, Clifford Meyer, Steve Miller, Matthew G. Milton, Tejaswini Mishra, Dianna E. Moore, Helen M. Moore, Jill E. Moore, Samuel H. Moore, Jennifer Moran, Ali Mortazavi, Jonathan M. Mudge, Nikhil Munshi, Rabi Murad, Richard M. Myers, Vivek Nandakumar, Preetha Nandi, Anil M. Narasimha, Aditi K. Narayanan, Hannah Naughton, Fabio C. P. Navarro, Patrick Navas, Jurijs Nazarovs, Jemma Nelson, Shane Neph, Fidencio Jun Neri, and The ENCODE Project Consortium. Expanded encyclopaedias of DNA elements in the human and mouse genomes. *Nature*, 583(7818):699–710, 2020.

[2] Mary A Allen, Hestia Mellert, Veronica Dengler, Zdenek Andryzik, Anna Guarnieri, Justin A Freeman, Xin Luo, William L Kraus, Robin D Dowell, and Joaquín M Espinosa. Global analysis of p53-regulated transcription identifies its direct targets and unexpected regulatory mechanisms. *eLife*, 3:e02200, 2014.

[3] Zdenek Andrysik, Matthew D. Galbraith, Anna L. Guarnieri, Sara Zaccara, Kelly D. Sullivan, Ahwan Pandey, Morgan MacBeth, Alberto Inga, and Joaquin M. Espinosa. Identification of a core TP53 transcriptional program with highly distributed tumor suppressive activity. *Genome Research*, 2017.

[4] Joseph G. Azofeifa, Mary A. Allen, Manuel E. Lladser, and Robin D. Dowell. An annotation agnostic algorithm for detecting nascent RNA transcripts in GRO-seq. 14(5):1070–1081.

[5] Felipe Beckedorff, Ezra Blumenthal, Lucas Ferreira daSilva, Yuki Aoi, Pradeep Reddy Cingaram, Jingyin Yue, Anda Zhang, Sadat Dokaneheifard, Monica Guiselle Valencia, Gabriel Gaidosh, Ali Shilatifard, and Ramin Shiekhattar. The human integrator complex facilitates transcriptional elongation by endonucleolytic cleavage of nascent transcripts. *Cell Reports*, 32(3), 2024/10/02 2020.

[6] Leighton J. Core, André L. Martins, Charles G. Danko, Colin T. Waters, Adam Siepel, and John T. Lis. Analysis of nascent RNA identifies a unified architecture of initiation regions at mammalian promoters and enhancers. *Nature Genetics*, 46(12):1311–1320, Dec 2014.

[7] Heather L. Drexler, Karine Choquet, and L. Stirling Churchman. Splicing kinetics and coordination revealed by direct nascent RNA sequencing through nanopores. *Molecular Cell*, 77(5):985–998.e8, 2020.

[8] Heather L. Drexler, Karine Choquet, Hope E. Merens, Paul S. Tang, Jared T. Simpson, and L. Stirling Churchman. Revealing nascent RNA processing dynamics with nano-COP. *Nature Protocols*, 16(3):1343–1375, 2021. Number: 3 Publisher: Nature Publishing Group.

[9] Sven Heinz, Christopher Benner, Nathanael Spann, Eric Bertolino, Yin C. Lin, Peter Laslo, Jason X. Cheng, Cornelis Murre, Harinder

Singh, and Christopher K. Glass. Simple combinations of lineage-determining transcription factors prime *cis*-regulatory elements required for macrophage and b cell identities. *Molecular Cell*, 38(4):576–589, 2010.

[10] Charity W. Law, Yunshun Chen, Wei Shi, and Gordon K. Smyth. voom: precision weights unlock linear model analysis tools for RNA-seq read counts. *Genome Biology*, 15(2):R29, 2014.

[11] Michael I. Love, Wolfgang Huber, and Simon Anders. Moderated estimation of fold change and dispersion for rna-seq data with deseq2. *Genome Biology*, 15(12):550, 2014.

[12] Zachary L Maas and Robin D Dowell. Internal and external normalization of nascent RNA sequencing run-on experiments. *BMC Bioinformatics*, 25(1):19, Jan 2024.

[13] Kirsten A. Reimer, Claudia A. Mimoso, Karen Adelman, and Karla M. Neugebauer. Co-transcriptional splicing regulates $3'$ end cleavage during mammalian erythropoiesis. *Molecular Cell*, 81(5):998–1012.e7, 2021.

[14] Mark D. Robinson, Davis J. McCarthy, and Gordon K. Smyth. edgeR: a bioconductor package for differential expression analysis of digital gene expression data. *Bioinformatics*, 26(1):139–140, 2010.

[15] Rutendo F. Sigauke, Lynn Sanford, Zachary L. Maas, Taylor Jones, Jacob T. Stanley, Hope A. Townsend, Mary A. Allen, and Robin D. Dowell. Atlas of nascent RNA transcripts reveals enhancer to gene linkages. *BMC Genomics*, in press, 2025.

[16] A. Vihervaara, D. B. Mahat, M. J. Guertin, T. Chu, C. G. Danko, J. T. Lis, and L. Sistonen. Transcriptional response to stress is pre-wired by promoter and enhancer architecture. *Nat Commun*, 8(1):255, 2017.

[17] Allen Wang, Feng Yue, Yan Li, Ruiyu Xie, Thomas Harper, Nisha A Patel, Kayla Muth, Jeffrey Palmer, Yunjiang Qiu, Jinzhao Wang, Dieter K Lam, Jeffrey C Raum, Doris A Stoffers, Bing Ren, and Maike Sander. Epigenetic priming of enhancers predicts developmental competence of hESC-derived endodermal lineage intermediates. *Cell Stem Cell*, 16(4):386–99, Apr 2015.

[18] Li Yao, Jin Liang, Abdullah Ozer, Alden King-Yung Leung, John T Lis, and Haiyuan Yu. A comparison of experimental assays and analytical methods for genome-wide identification of active enhancers. *Nature Biotechnology*, pages 1–10, 2022.
